# Identifying the cognitive processes underpinning hippocampal-dependent tasks

**DOI:** 10.1101/377408

**Authors:** Ian A. Clark, Victoria Hotchin, Anna Monk, Gloria Pizzamiglio, Alice Liefgreen, Eleanor A. Maguire

## Abstract

Autobiographical memory, future thinking and spatial navigation are critical cognitive functions that are thought to be related, and are known to depend upon a brain structure called the hippocampus. Surprisingly, direct evidence for their interrelatedness is lacking, as is an understanding of why they might be related. There is debate about whether they are linked by an underlying memory-related process or, as has more recently been suggested, because they each require the endogenous construction of scene imagery. Here, using a large sample of participants and multiple cognitive tests with a wide spread of individual differences in performance, we found that these functions are indeed related. Mediation analyses further showed that scene construction, and not memory, mediated (explained) the relationships between the functions. These findings offer a fresh perspective on autobiographical memory, future thinking, navigation, and also on the hippocampus, where scene imagery appears to play an influential role.

---

Our past experiences are captured in autobiographical memories that serve to sustain our sense of self, enable independent living and prolong survival (Tulving, 2002). Consequently, a key aim of cognitive psychology and neuropsychology has been to understand how such memories are formed and recollected. There is wide agreement that a brain structure called the hippocampus plays a key role in supporting autobiographical memories. Patients with hippocampal damage are impaired at recalling past experiences (Scoville & Milner, 1957; see also Clark & Maguire, 2016; Verfaellie & Keane, 2017; Winocur & Moscovitch, 2011), and the hippocampus is consistently engaged during functional MRI studies of autobiographical memory retrieval (Cabeza & St. Jacques, 2007; Svoboda, McKinnon, & Levine, 2006). Consequently, the hippocampus and autobiographical memory have become synonymous.

However, the hippocampus has been associated with functions beyond autobiographical memory. The animal literature has, for many years, placed spatial navigation at the heart of hippocampal processing (Moser, Kropff, & Moser, 2008; O’Keefe & Dostrovsky, 1971; O’Keefe & Nadel, 1978), with concordant findings in humans (Ekstrom et al., 2003; Epstein, Patai, Julian, & Spiers, 2017; Maguire et al., 2000). Work over the past decade has also linked the hippocampus with thinking about the future (Addis, Wong, & Schacter, 2007; Hassabis, Kumaran, Vann, & Maguire, 2007), the imagination of scenes and events (Hassabis, Kumaran, Vann, et al., 2007; Schacter et al., 2012), the perception of scenes (Graham, Barense, & Lee, 2010; McCormick, Rosenthal, Miller, & Maguire, 2017) and specific aspects of visuospatial processing, including perceptual richness, a sense of reliving and imagery content (Andrews-Hanna, Reidler, Sepulcre, Poulin, & Buckner, 2010; St-Laurent, Moscovitch, & McAndrews, 2016; St. Jacques, Conway, Lowder, & Cabeza, 2010).

The link between autobiographical memory, the construction of scene imagery in the imagination (scene construction) and thinking about the future has come under increasing scrutiny. Studies of amnesic patients have reported deficits in tasks assessing each of these functions (Hassabis, Kumaran, Vann, et al., 2007; Klein, Loftus, & Kihlstrom, 2002; Rosenbaum et al., 2005; Tulving, 1985). In neuroimaging studies, the recruitment of the same neural network, including the hippocampus, has been observed when thinking about the past, the future or atemporal events and scenes with no obvious focus in time (Buckner & Carroll, 2007; Hassabis & Maguire, 2007; Schacter, Addis, & Buckner, 2007). In addition, comparisons of behavioural measures have highlighted similarities in terms of ratings of vividness and the amount and type of details for past, future and atemporal events (D’Argembeau & Van der Linden, 2006; de Vito, Gamboz, & Brandimonte, 2012). Overall, therefore, autobiographical memory, scene construction and thinking about the future seem to involve the hippocampus, with parallels also in the pattern of behavioural outcomes. Yet, conceptually, they are different processes not least in terms of the temporal context within which the scene or event is imagined. The question, therefore, arises as to what does the hippocampus do in the service of each of these functions?

One suggestion is that autobiographical memory provides the building blocks for thinking about the future and imagining atemporal scenes and events and, as such, their dependence on the hippocampus is fundamentally mnemonic (Moscovitch, Cabeza, Winocur, & Nadel, 2016; Schacter et al., 2012; Sheldon & Levine, 2016). This is based upon the suggestion that autobiographical memory recall is a constructive process that recombines different elements to recreate memories (e.g. Schacter et al., 2012). This information is also available for the construction of non-autobiographical memory events. In this regard the autobiographical memory system is equally well equipped to imagine future or atemporal events as well as recalling the past (see also St. Jacques, Carpenter, Szpunar, & Schacter, 2018; Thakral, Benoit, & Schacter, 2017).

An alternative view is that the mental construction of scene imagery is a key process that autobiographical memory, future thinking and spatial navigation have in common (Maguire & Mullally, 2013; see also, Robin, 2018; Rubin & Umanath, 2015 for related theoretical viewpoints). A scene is a naturalistic three-dimensional spatially coherent representation of the world typically populated by objects and viewed from an egocentric perspective. When most people recall the past, imagine the future or plan a route during navigation, scenes feature prominently. An individual’s ability to use scene imagery, or spatial context, to imagine or recall an event, has been shown to predict the vividness and detail of the imagined scenario (Arnold, McDermott, & Szpunar, 2011; D’Argembeau & Van der Linden, 2004; Hebscher, Levine, & Gilboa, 2017; Robin & Moscovitch, 2014; Robin, Wynn, & Moscovitch, 2016; Sheldon & Chu, 2017; Szpunar & McDermott, 2008). Furthermore, damage limited to the hippocampus is known to impede the ability to construct endogenous scene imagery (Andelman, Hoofien, Goldberg, Aizenstein, & Neufeld, 2010; Hassabis, Kumaran, Vann, et al., 2007; Maguire & Mullally, 2013; Race, Keane, & Verfaellie, 2011; Rosenbaum, Gilboa, Levine, Winocur, & Moscovitch, 2009). The mental construction of scenes is, therefore, both reliant upon hippocampal functionality and related to autobiographical memory, future thinking and spatial navigation.

There is, however, a dearth of evidence available that permits adjudication between a mnemonic or scene construction account of hippocampal function. Arguably, extant evidence highlights the importance of scene construction over autobiographical memory (de Vito et al., 2012; Palombo, Hayes, Peterson, Keane, & Verfaellie, 2018; Robin & Moscovitch, 2014; but see also, Addis, Cheng, Roberts, & Schacter, 2011; Roberts, Schacter, & Addis, 2017). To the best of our knowledge, there are no large scale individual differences studies systematically examining the direct relationships between scene construction, autobiographical memory, thinking about the future and spatial navigation.

We, therefore, had two overarching goals in the current study. First, we sought to investigate whether scores on tasks assessing scene construction, autobiographical memory, future thinking and spatial navigation were related. Second, if they were related, we wished to unpack these relationships further by attempting to pinpoint whether the link between them could be best explained by either a scene construction or an autobiographical memory process. Given that the scene construction deficit of hippocampal-damaged patients is evident even on non-mnemonic tasks, for example, the visual perception of scenes (Lee et al., 2005; McCormick et al., 2017), we hypothesised that the relationship, if any, between the tasks would be best explained by scene construction rather than by autobiographical memory.

To address our first goal, we conducted a principal component analysis (PCA) involving a large range of cognitive tests. This allowed us to assess whether or not performance on tasks examining scene construction, autobiographical memory, future thinking and navigation was related in the presence of other cognitive tasks.

To pursue our second goal, we performed a series of mediation analyses. This approach allowed us to investigate how scores on the tasks of interest were related. Mediation analyses focus on whether one variable can explain (mediate) the relationship between two different variables. In short, we expected that if the cognitive process linking the different tasks together was related to scenes, then scene construction performance would mediate the task relationships. By contrast, if the underlying process was related to autobiographical memory, then autobiographical memory performance would be found to mediate.

We recruited a large group of participants and assessed their performance on a comprehensive battery of cognitive tasks, including measures of scene construction, autobiographical memory, future thinking and navigation. Tasks were chosen from the published literature because of their confirmed reliance (or non-reliance) upon the hippocampus.

## Method

### Participants

Two hundred and seventeen individuals were recruited. They were aged between 20 and 41 years old, had English as their first language and reported no psychological, psychiatric, neurological or behavioural health conditions. The age range was restricted to 20-41 to limit the possible effects of ageing. Participants reporting hobbies or vocations known to be associated with the hippocampus (e.g. licensed London taxi drivers) were excluded. The mean age of the sample was 29.0 years (95% CI; 20, 38) and included 109 females and 108 males. Participants were reimbursed £10 per hour for taking part which was paid at study completion. All participants gave written informed consent and the study was approved by the University College London Research Ethics Committee. APA ethical standards were complied with in regards to the treatment of the participants.

The sample size was determined at 216 during study design to be robust to employing different statistical approaches when answering multiple questions of interest. Specifically, the sample allows for sufficient power to identify medium effect sizes when conducting regression analyses, which form the basis of mediation analyses, at alpha levels of .01 (Cohen, 1992). Importantly, the sample size is also large enough to conduct mediation analyses and structural equation modelling (Anderson & Gerbing, 1988). A final sample of 217 was obtained due to over recruitment.

### Procedure

Participants completed the study over three separate visits. The order of tests within each visit was the same for all participants (see Task Order). Task order was arranged so as to avoid task interference, for example, not having a verbal test followed by another verbal test, and to provide sessions of approximately equal length (~3-3.5 hours, including breaks). All participants completed all parts of the study.

### Cognitive Tests

#### Measures of primary interest

Our main interest was in scene construction, autobiographical memory, future thinking and navigation; tasks which are known to recruit or require the hippocampus in order to be successfully completed. All tasks are published and were performed and scored as per their published use. Given the extensive task battery that was used, only the main outcome measure was used for each task in order to reduce potential issues surrounding multiple comparisons and false positives. Here, for the reader’s convenience, we describe each task briefly.

##### Scene construction test

(Hassabis, Kumaran, Vann, et al., 2007). This test measures a participant’s ability to mentally construct a visual scene. Participants construct different scenes of commonplace settings. For each scene, a short cue is provided (e.g. imagine lying on a beach in a beautiful tropical bay) and the participant is asked to imagine the scene that is evoked and then describe it out loud in as much detail as possible. Recordings are transcribed for later scoring. Participants are explicitly told not to describe a memory, but to create a new scene that they have never experienced before.

The overall outcome measure is an “experiential index” which is calculated for each scene and then averaged. In brief, it is composed of four elements: the content, participant ratings of their sense of presence (how much they felt like they were really there) and perceived vividness, participant ratings of the spatial coherence of the scene, and an experimenter rating of the overall quality of the scene.

Double scoring was performed on 20% of the data. We took the most stringent approach to identifying across-experimenter agreement. Inter-class correlation coefficients, with a two way random effects model looking for absolute agreement indicated excellent agreement among the experimenter ratings (minimum score of .9; see Supplementary Materials, Table S1). For reference, a score of .8 or above is considered excellent agreement beyond chance.

##### Autobiographical Interview

(AI; Levine, Svoboda, Hay, Winocur, & Moscovitch, 2002). In the AI participants are asked to provide autobiographical memories from a specific time and place over four time periods – early childhood (up to age 11), teenage years (aged from 11-17), adulthood (from age 18 years to 12 months prior to the interview; two memories are requested) and the last year (a memory from the last 12 months). Recordings are transcribed for later scoring.

In contrast to the other tasks, the AI has two main outcome measures, both of which are consistently reported in the literature. Memories are scored to collect “internal” and “external” details of the event. Importantly, these two scores represent different aspects of autobiographical memory recall. Internal details are those describing the event in question (i.e. episodic details). External details describe semantic information concerning the event, or non-event information. Internal events are therefore thought to be hippocampal-dependent, while external events are not. As such, in line with the published literature, we report both outcome measures. The two AI scores are obtained by separately averaging performance for the internal and external details across five autobiographical memories. Our double scoring produced excellent agreement across the experimenters (minimum score of .81; see Supplementary Materials, Table S2).

##### Future thinking test

(Hassabis, Kumaran, Vann, et al., 2007). This test follows the same procedure as the scene construction test, but requires participants to imagine three plausible future scenes involving themselves (an event at the weekend; next Christmas; the next time they will meet a friend). Participants are explicitly told not to describe a memory, but to create a new future scene. Recordings are transcribed for later scoring. The scoring procedures are the same as for scene construction. Double scoring identified excellent agreement across the experimenters (minimum score of .88; see Supplementary Materials, Table S3).

##### Navigation tests

(Woollett & Maguire, 2010). Navigation ability is assessed using movies of navigation through an unfamiliar town. Movie clips of two overlapping routes through this real town (Blackrock, in Dublin, Ireland) are shown to participants four times.

Five tasks are used to assess navigational ability. First, following each viewing of the route movies, participants are shown four short clips – two from the actual routes, and two distractors. Participants indicate whether they have seen each clip or not. Second, after all four route viewings are completed, recognition memory for scenes from the routes is tested. A third test involves assessing knowledge of the spatial relationships between landmarks from the routes. Fourth, route knowledge is examined by having participants place photographs from the routes in the correct order as if travelling through the town. Finally, participants draw a sketch map of the two routes including as many landmarks as they can remember. Sketch maps are scored in terms of the number of road segments, road junctions, correct landmarks, landmark positions, the orientation of the routes and an overall map quality score from the experimenters. Double scoring was performed on 20% of the sketch maps finding excellent agreement (minimum of .89; see Supplementary Materials, Table S4). An overall navigation score is calculated by combining scores from all of the above tasks.

#### Additional measures

We administered a range of other tasks to participants which enabled us to further profile their cognition. In brief, estimates of IQ were obtained using the Test of Premorbid Functioning (TOPF; Wechsler, 2011). The number of correct responses was converted to an estimate of Full Scale IQ (FSIQ) as per the TOPF scoring procedure. General intellect and executive functioning were measured using the Matrix Reasoning subtest of the Wechsler Adult Intelligence Scale IV (WAIS-IV; Wechsler, 2008), the Brixton Spatial Anticipation Test (Burgess & Shallice, 1997) and the F-A-S verbal fluency task (F-A-S; Strauss, Sherman, & Spreen, 2006). Working memory/attention was assessed using the Digit Span subtest of the WAIS-IV and the Symbol Span subtest of the Wechsler Memory Scale IV (WMS-IV; Wechsler, 2009).

Visuospatial recall was examined using the Rey–Osterrieth Complex Figure (ROCF; Rey, 1941). In addition, we also used an object-place association test which required participants to learn the locations of 16 objects presented simultaneously on a white computer screen (adapted from Woollett & Maguire, 2009). The outcome measure was the number of trials (maximum of 6) taken to correctly learn the location of all the objects, with a score of 7 if the array was never learnt (this was reverse scored for ease of interpretation with the other tasks).

Verbal recall was assessed using the Rey Auditory Verbal Learning Test (RAVLT; see Strauss et al., 2006), and the Logical Memory and Verbal Paired Associates subtests of the WMS-IV (Wechsler, 2009). Two additional verbal recall tasks were also included (Clark, Kim, & Maguire, 2018). A limitation of the WMS Verbal Paired Associates task is its reliance on concrete, imageable words (Clark & Maguire, 2016; Maguire & Mullally, 2013). We therefore included two additional versions of this task. In one case, only concrete, imageable words are used while the other comprises only abstract, non-imageable words. The two tests are precisely matched apart from the imageability of the words. For all of these recall tasks, the delayed recall scores were used as our primary data as they are most sensitive to hippocampal damage (Squire, 1992).

Recognition memory was assessed using the Warrington Recognition Memory Tests for words, faces and scenes (Cipolotti & Maguire, 2003; Warrington, 1984). Semantic memory was assessed using the “Dead or Alive” task which probes general knowledge about whether famous individuals have died or are still alive (Kapur, Young, Bateman, & Kennedy, 1989).

General visuospatial processing was assessed using the Paper Folding test (Ekstrom, French, Harman, & Dermen, 1976) which measures a participant’s ability to transform images of spatial patterns into different arrangements. Perceptual processing was assessed using scene description and boundary extension tasks (Mullally, Intraub, & Maguire, 2012). The scene description task requires participants to describe a picture of a scene. The content of participants’ descriptions is scored across a number of categories and summed to provide a total content score. Double scoring was performed on 20% of the descriptions finding excellent agreement (minimum of .85; see Supplementary Materials, Table S5). Boundary extension occurs when individuals who are viewing scenes automatically imagine what might be beyond the view, and consequently later misremember having seen a greater expanse of the scene (Intraub & Richardson, 1989). To test this, participants are briefly presented on each trial with two pictures in rapid succession and are asked to rate whether the second picture is of a closer perspective (when boundary extension is induced), exactly the same (the correct answer), or further away. Unbeknownst to participants, the majority of images are exactly the same. The outcome measure was the proportion of same trials classed as closer-up.

#### Task order

The tests were conducted in the following order: In session 1 – Concrete Verbal Paired Associates (learning), Warrington Recognition Memory Test for scenes, Dead or Alive task, Symbol Span test, scene description task, Concrete Verbal Paired Associates (delayed recall), Logical Memory test (learning), ROCF (copy), TOPF, Warrington Recognition Memory Test for faces, Brixton Spatial Anticipation Test, Logical Memory test (delayed recall), ROCF (delayed recall), Warrington Recognition Memory Test for words. In session 2 – navigation tests, followed by Abstract Verbal Paired Associates (learning, with delayed recall 30 minutes later). In session 3 – scene construction test, future thinking test, RAVLT (learning), Paper Folding test, Digit Span test, Matrix Reasoning, RAVLT (delayed recall), Autobiographical Interview, WMS Verbal Paired Associates (learning), object-place association test, boundary extension task, WMS Verbal Paired Associates (delayed recall), F-A-S task.

### Statistical analyses

Data are summarised using means and 95% confidence intervals, calculated in SPSS v22. PCA was performed using SPSS v22, with varimax rotation and a cut-off at an eigenvalue of 1. Regression analyses with standardised beta values and confidence intervals were performed in R v3.4. Mediation and sensitivity analyses were performed using the R Causal Mediation Analysis package v4.4.6 (Imai, Keele, & Tingley, 2010). Structural Equation Modelling (SEM) was performed using the R Lavaan package v0.6-1.1178 (Rosseel, 2012) and assessed for model fit as per the criteria of Hu and Bentler (1999). Effect sizes are reported as R^2^ values for regressions, including those regressions used in the mediation and SEM analyses (adjusted R^2^ when multiple variables were included) and as sensitivity analyses for the mediation analyses. There were no missing data, and no data needed to be removed from any analysis.

## Results

A summary of the outcome measures for the cognitive tasks is presented in Table 1. A wide range of scores was obtained for all variables.

**Table 1.** Means with 95% confidence intervals (CI) for all cognitive tasks

| Variable | Mean | Lower CI | Upper CI |
| --- | --- | --- | --- |
| Scene Construction Experiential Index (/60) | 40.50 | 29.50 | 50.13 |
| Autobiographical Memory Internal Details (total number) | 23.95 | 13.80 | 37.41 |
| Autobiographical Memory External Details (total number) | 5.35 | 1.40 | 11.24 |
| Future Thinking Experiential Index (/60) | 39.12 | 25.00 | 49.99 |
| Navigation (/250) | 143.46 | 88.90 | 201.50 |
| Full Scale Intelligence Quotient* | 102.75 | 92.04 | 114.35 |
| Matrix Reasoning scaled score (/19) | 12.53 | 8.00 | 17.00 |
| Brixton Spatial Anticipation Test scaled score (/10) | 7.87 | 5.00 | 10.00 |
| F-A-S Verbal Fluency (total number of words) | 49.09 | 30.90 | 69.00 |
| Digit Span scaled score (/37 – sum of backwards and forwards) | 22.08 | 14.00 | 34.00 |
| Symbol Span scaled score (/19) | 9.35 | 6.00 | 13.00 |
| Rey-Osterrieth Complex Figure delayed recall (/36) | 22.28 | 12.45 | 31.00 |
| Object-Place Association Test (/6) | 2.31 | 0.00 | 6.00 |
| Rey Auditory Verbal Learning Test delayed recall (/15) | 12.92 | 8.90 | 15.00 |
| Logical Memory delayed recall scaled score (/19) | 12.58 | 8.00 | 17.00 |
| WMS Verbal Paired Associates delayed recall (/14) | 13.38 | 10.00 | 14.00 |
| Concrete Verbal Paired Associates delayed recall (/14) | 12.94 | 8.00 | 14.00 |
| Abstract Verbal Paired Associates delayed recall (/14) | 7.03 | 1.00 | 13.10 |
| Warrington Recognition Memory Test for Words scaled score (/15) | 12.75 | 7.00 | 14.00 |
| Warrington Recognition Memory Test for Faces, scaled score (/18) | 11.00 | 4.00 | 16.00 |
| Warrington Recognition Memory Test for Scenes raw score (/50) | 43.35 | 35.00 | 49.00 |
| Dead or Alive Task proportion correct (%) | 81.32 | 66.12 | 94.52 |
| Paper Folding Test (/20) | 13.14 | 6.00 | 19.00 |
| Scene Description (total number of details) | 24.88 | 15.90 | 37.10 |
| Boundary Extension proportion of ‘closer’ responses (%) | 42.26 | 8.33 | 79.17 |

*Note.* Task order is for display purposes only. \*Estimated from the Test of Premorbid Functioning. WMS = Wechsler Memory Scale.

### How are the tasks interrelated?

We first asked whether performance across the tasks of primary interest (scene construction, autobiographical memory, future thinking and navigation), was related. If, in line with our prediction, these tasks share an underlying cognitive process, then performance on one task should be related to performance on the others. More generally, we also sought to investigate this within the wider context of the other cognitive tasks.

We performed a PCA using all of the tasks in order to avoid selection bias – we had no reason to exclude any task from the PCA. Varimax rotation was applied (to allow for cross-over between the derived components) and the minimum eigenvalue was set to 1. The PCA identified 7 components that explained 59.24% of the variance. Examination of the Scree plot supported the 7 component solution (see Supplementary Materials and Figure S1).

Naming of the components was determined by the tasks that most strongly loaded on to each (see Table 2 for the proportion of variance explained by each component, Table 3 for the tasks in each component and their weightings and Supplementary Materials Table S6 for all weightings). Component 1 comprised tasks with a particularly strong spatial component (e.g. navigation, object-place association, Paper Folding). Notably, this was regardless of whether or not memory was required; for example, the Paper Folding test and the Brixton Spatial Anticipation Test had minimal memory requirements. Component 2 contained all of the verbal memory tasks. Component 3 comprised those tasks typically thought to assess general IQ or executive function. Matrix Reasoning and the estimate of FSIQ from the TOPF are designed to be measures of general IQ (Wechsler, 2008, 2011), the Symbol Span and Digit Span tests measure executive function, working memory and attention (Wechsler, 2008, 2009), and the F-A-S is reported as an executive function task (Strauss, Sherman, & Spreen, 2006). Finally, while the Abstract Verbal Paired Associates test is a verbal memory task (and aligns also with component 2), it is a more challenging task than the other verbal memory tasks – as shown by the performance scores in Table 1 – and has also been found to require the recruitment of frontal ‘executive’ brain regions (Clark et al. 2018), suggesting that processing of Abstract Verbal Paired Associates may reflect general IQ as well as verbal memory. Component 4 involved three of our tasks of primary interest - scene construction, autobiographical memory (internal details) and future thinking, and the inclusion of the simple scene description task (which also loaded onto the perceptual component). For convenience, we refer to this component as the ‘Scene’ component, as per our hypothesis that these tasks have scenes in common, but acknowledge that this remains to be tested in our following analyses. Component 5 contained the three recognition memory tests and component 6 the two semantic tasks. Finally, component 7, while also using scene-based stimuli, contained the two tasks that primarily assessed visual perception.

There are, however, two potential limitations to the PCA that should be noted. First, it could be suggested that inclusion of FSIQ in the PCA reduces the pool of domain-specific variance that can be partitioned because FSIQ could be regarded as a more non-specific cognitive task. To ensure FSIQ scores were not biasing the results, we performed the PCA again without the FSIQ scores, finding a near identical pattern of results (see Supplementary Materials, Table S7). The only notable effect of removing FSIQ was that components 3 and 4 were switched, with the Scene component now explaining a greater proportion of the variance than the IQ/Executive Function component.

Second, the selection of the number of components from a PCA is, at least in part, a subjective decision. From the scree plot, it could be argued that our PCA should only result in four factors, rather than seven. However, as has been widely discussed in the literature, determining the number of components to include requires the balancing of parsimony and plausibility (see, for example, Fabrigar, Wegener, MacCallum, & Strahan, 1999), with the selection of too few components being typically regarded as a much more severe error than too many (Cattell, 1978; Fava & Velicer, 1992; Rummel, 1970; Wood, Tataryn, & Gorsuch, 1996). As detailed in the Supplementary Materials, restricting the model to four components meant that two tasks (Dead or Alive, Boundary Extension) loaded onto none of the components. Furthermore, limiting the solution to four components led to the loss of the Recognition Memory, Semantic Memory and Perception components, resulting, for example, in the Warrington Recognition Memory Test for faces loading onto the Verbal Memory component. As such, specifying only four components obscured theoretically plausible and relevant components as well as leading to difficulties in interpretation. On the other hand, the seven component solution was statistically valid, created a clear factor structure with all the tests included, and all components had both statistical and theoretical value.

We also note that our main focus here was on scene construction, autobiographical memory, future thinking and navigation, and investigating how performance on these tasks is related in the presence of other cognitive tasks. Regardless of the above issues – the inclusion/exclusion of FSIQ and the selection of either four or seven components – our main tasks of interest followed the same loadings and the Scene and Spatial components remained within the top four explanatory components.

**Table 2.** Proportion of variance explained by each PCA component

| PCA Component | Variance Explained |
| --- | --- |
| Total | 59.24% |
| C1 – Spatial Processing | 12.44% |
| C2 – Verbal Memory | 10.64% |
| C3 – General IQ/Executive Function | 9.85% |
| C4 – Scenes | 9.23% |
| C5 – Recognition Memory | 6.81% |
| C6 – Semantic Memory | 5.28% |
| C7 – Perception | 5.01% |

**Table 3.** Details of the Principal Component Analysis with varimax rotation of the cognitive tasks

| Cognitive Task | Spatial | Verbal | IQ/Executive Function | Scenes | Recognition Memory | Semantic Memory | Perception |
| --- | --- | --- | --- | --- | --- | --- | --- |
| Rey-Osterrieth Complex Figure delayed recall | .72 |  |  |  |  |  |  |
| Paper Folding Test | .72 |  |  |  |  |  |  |
| Navigation | .66 |  |  |  |  |  |  |
| Object-Place Association Test | .65 |  |  |  |  |  |  |
| Brixton Spatial Anticipation Test | .42 |  |  |  |  |  |  |
| Warrington Recognition Memory Test for Scenes | .54 |  |  |  | .41 |  |  |
| Matrix Reasoning | .51 |  | .49 |  |  |  |  |
| Symbol Span | .46 |  | .46 |  |  |  |  |
| Rey Auditory Verbal Learning Test delayed recall |  | .74 |  |  |  |  |  |
| Concrete Verbal Paired Associates delayed recall |  | .67 |  |  |  |  |  |
| Logical Memory delayed recall |  | .66 |  |  |  |  |  |
| WMS Verbal Paired Associates delayed recall |  | .62 |  |  |  |  |  |
| Abstract Verbal Paired Associates delayed recall |  | .61 | .46 |  |  |  |  |
| Digit Span |  |  | .74 |  |  |  |  |
| Full Scale Intelligence Quotient |  |  | .68 |  |  |  |  |
| F-A-S Verbal Fluency |  |  | .62 |  |  |  |  |
| Scene Construction Experiential Index |  |  |  | .87 |  |  |  |
| Future Thinking Experiential Index |  |  |  | .85 |  |  |  |
| Autobiographical Memory Internal Details |  |  |  | .62 |  |  |  |
| Scene Description |  |  |  | .37 |  |  | .50 |
| Warrington Recognition Memory Test for Words |  |  |  |  | .67 |  |  |
| Warrington Recognition Memory Test for Faces |  |  |  |  | .79 |  |  |
| Autobiographical Memory External Details |  |  |  |  |  | .69 |  |
| Dead or Alive Task |  |  |  |  |  | .62 |  |
| Boundary Extension |  |  |  |  |  |  | .84 |

*Note.* Task order is for display purposes only. Only values over .35 are reported for ease of viewing, full results are presented in Supplementary Materials Table S6. WMS = Wechsler Memory Scale.

In summary, performance on the scene construction, autobiographical memory (internal details) and future thinking tasks all aligned onto the same component. This demonstrates that these tasks are strongly related in cognitive terms. However, surprisingly, the navigation task did not load onto this component, but instead loaded onto the spatial component – a point we will return to later.

While the PCA can tell us about the main relationships between tasks, it cannot inform about the nature of the underlying processes. We therefore proceeded to perform additional analyses to examine this.

### What cognitive process(es) underpin the Scene component?

The PCA analysis identified that, with the exception of navigation, our tasks of primary interest – scene construction, autobiographical memory internal details (henceforth referred to as autobiographical memory) and future thinking – all loaded onto one component – Scenes. As the scene description task also loaded onto the Perception component as well as the Scene component, it was not included in the following analyses to allow for the assessment of just the pure elements of the Scene component.

We used mediation analyses to investigate possible processes underpinning the Scene component. This method aims to explain the mechanisms and/or processes underlying the relationship between two variables via the inclusion of a third variable. If the third variable fully mediates the original relationship, this provides evidence that the link between the original variables can be explained solely due to the mediating variable. This is known as an indirect effect. On the other hand, if no indirect effect is identified, leaving only the direct relationship between the original variables, it can be concluded that the mediating variable is not involved in the original relationship. For a mediation analysis to be possible, there are two main requirements, as described by Baron & Kenny (1986). First, the independent variable must be a predictor of the dependent variable. Second, the independent variable must predict the mediator variable. The first requirement has, however, been further scrutinised, with some suggesting that an initial direct relationship between the independent variable and the dependent variable is not required when there is a strong a priori belief that the effect size is small or suppression is a possibility (e.g. Shrout & Bolger, 2002). However, given our substantial sample size and ability to detect small effect sizes, we followed the more stringent requirements set out by Baron & Kenny (1986) to reduce the possibility of false positives. A mediation analysis then looks at the difference between predicting the dependent variable from just the independent variable, in comparison to predicting the dependent variable from the independent variable *and* the mediator variable. If the relationship between the independent and dependent variable is reduced, or lost, with the inclusion of the mediator, an indirect effect has occurred.

Mediation can, therefore, be applied to our question in the following manner – if the process linking the Scene component tasks is, as we hypothesise, related to scenes, the scene construction task should mediate the relationship between autobiographical memory and future thinking. Alternatively, if, as hypothesised by others, the underlying process is associated with autobiographical memory, then autobiographical memory will mediate the relationship between scene construction and future thinking. This was, therefore, our first analysis.

Before reporting the results, it is worth explaining the presentation format. A mediation analysis has two main steps. First, the initial regressions are performed to ensure mediation is possible. For ease of reading, the full details of each individual regression are reported in the Supplementary Materials and just the unstandardised coefficients are reported in the main body of the text. Second, the mediation analysis itself is performed. The mediation analysis provides two outcome measures: (1) the mediation, or indirect, effect; if this is significant there is an effect of mediation, and (2) the direct effect; if this is significant then a relationship remains between the original variables even with the inclusion of the mediator. If just the indirect effect is significant, then a full mediation has occurred. This means that all of the relationship between the independent and dependent variable can be explained by the mediator. If both the indirect and the direct effect are significant then a partial mediation has occurred. This means that some of the relationship between the independent and dependent variables can be explained by the mediator, but the independent variable still contributes to the relationship. If only the direct effect is significant, then mediation has not occurred.

In a similar manner to other statistical analyses, it is important to look not just at whether a result is significant, but how robust this effect is. For mediation, this is done via sensitivity analyses. Sensitivity analyses test how well the (indirect or direct) effect holds if additional variance is introduced into the necessary assumptions made to perform the analysis (see Imai et al., 2010). Sensitivity analyses are different from effect sizes in that there are no specific cut offs. Instead, they are used comparatively. As such, sensitivity is reported here in two forms, first, as a single value (between −1 and 1) which represents the amount of additional variance needed to reduce the effect seen to 0. A higher absolute value represents a more robust effect. Second, we also display sensitivity as a plot showing the effect of varying the additional variance on the indirect or direct effect. This allows for a visual interpretation of the robustness of the effect.

Returning to the analyses, our overarching question was whether a scene construction or an autobiographical memory process best explained the relationships identified between scene construction, autobiographical memory and future thinking. We, therefore, systematically examined the different combinations of the relationships between these three variables. First, we investigated whether scene construction mediated the relationship between autobiographical memory and future thinking, or whether autobiographical memory mediated between scene construction and future thinking. Second, we also examined these relationships when future thinking was included as the independent instead of the dependent variable. Finally, for completeness, we investigated whether future thinking mediated the relationship between scene construction and autobiographical memory. Testing each of these in turn resulted in a complete examination of possible mediations between the three tasks.

First, therefore, we sought to examine whether scene construction mediated the relationship between autobiographical memory and future thinking, and whether autobiographical memory mediated the relationship between scene construction and future thinking. The results of the mediation analyses are shown in Table 4 and Figure 1 (see Supplementary Materials, Table S8 for the full break down of each individual regression). Figure 1a shows the relationship between autobiographical memory and future thinking, mediated by scene construction. As expected, autobiographical memory alone was associated with both future thinking (Beta = .39, *p* < .001) and scene construction (Beta = .36, *p* < .001). This shows that mediation by scene construction was possible. Indeed, with the inclusion of scene construction as a mediator, autobiographical memory was no longer related to future thinking (Beta = .063, *p* = .18), while scene construction was (Beta = .90, *p* < .001). Mediation analysis revealed a significant indirect effect of scene construction, with no direct effect of autobiographical memory (Table 4a). This, therefore, suggests that scene construction fully mediated (explained) the relationship between autobiographical memory and future thinking.

**Table 4.** Mediation analyses of the Scene component variables when future thinking is the dependent variable

|  | **Beta (95% CI)** | ***p*** | **Sensitivity (*p*)** |
| --- | --- | --- | --- |
| **a** |  |  |  |
| **Autobiographical Memory to Future Thinking, mediated by Scene Construction** |  |  |  |
| Indirect Effect | .32 (.23, .43) | < .001 | .75 |
| Direct Effect | .06 (-.03, .15) | .17 | -.2 |
| Total | .39 (.26, .51) | < .001 | n/a |
| **b** |  |  |  |
| **Scene Construction to Future Thinking, mediated by Autobiographical Memory** |  |  |  |
| Indirect Effect | .032 (-.014, .08) | .16 | .1 |
| Direct Effect | .90 (.80, 1.02) | < .001 | -.95 |
| Total | .94 (.84, 1.04) | < .001 | n/a |

**Figure 1.**
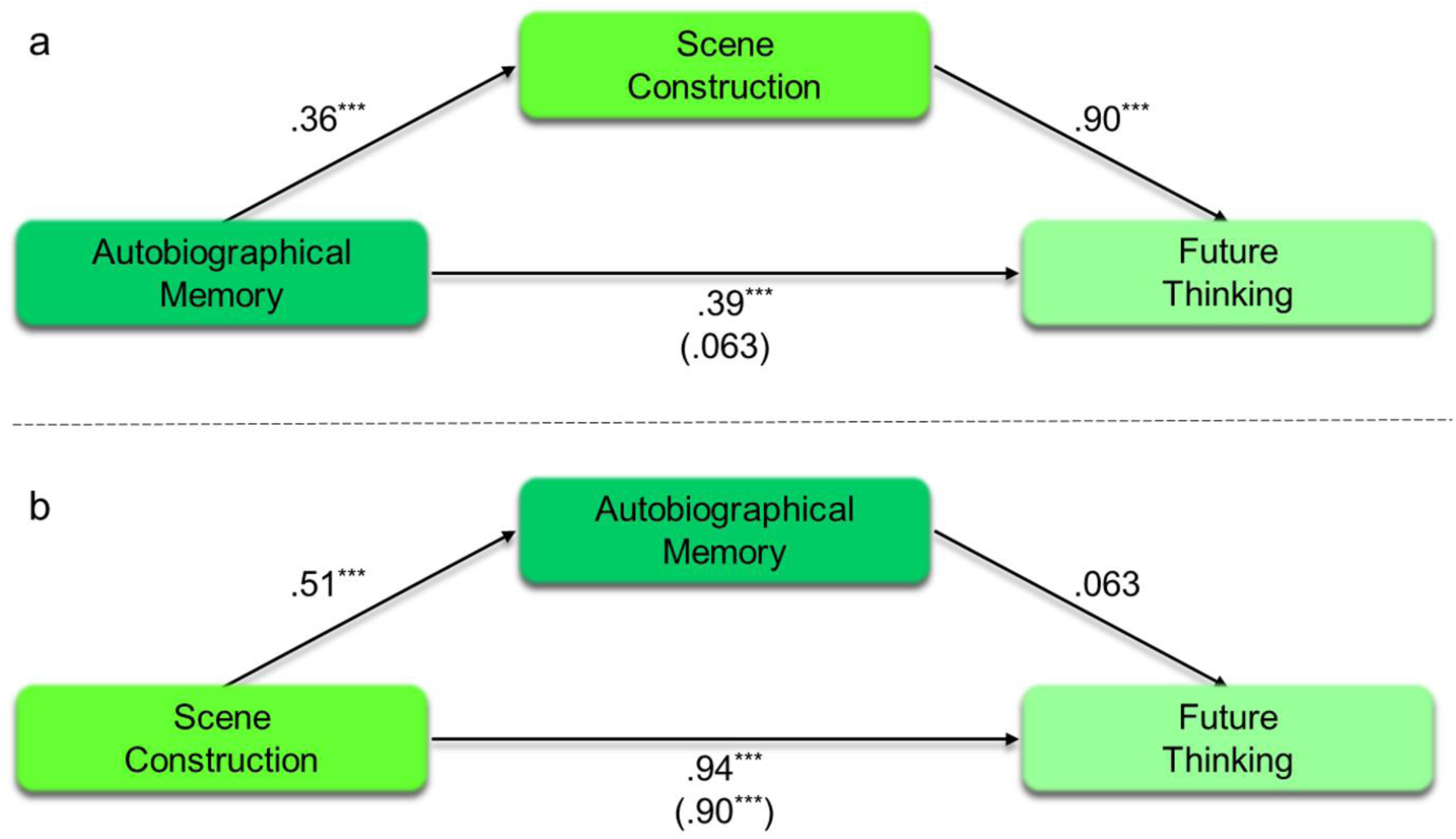
Mediation analyses of the Scene component variables when future thinking is the dependent variable. (a) Autobiographical memory to future thinking, mediated by scene construction. (b) Scene construction to future thinking, mediated by autobiographical memory. The numbers in brackets show the effect of the independent variable on the dependent when the mediation variable was also taken into consideration. ^***^*p* < .001.

Table 4b and Figure 1b show the equivalent analysis where autobiographical memory was placed as the mediator between scene construction and future thinking. As would be expected, the result matches the previous analysis, but with the indirect and direct effects switched. As with autobiographical memory, scene construction alone was associated with both future thinking (Beta = .94, *p* < .001) and autobiographical memory (Beta = .51, *p* < .001). This means that mediation by autobiographical memory was possible. However, including autobiographical memory as the mediator failed to show a relationship between autobiographical memory and future thinking (Beta = .063, *p* = .18), while the relationship between scene construction and future thinking remained significant (Beta = .90, *p* < .001). This was confirmed by the mediation analysis finding no indirect effect of autobiographical memory in comparison to the significant direct effect of scene construction. In other words, autobiographical memory could not explain the relationship between scene construction and future thinking. This is in contrast to the previous analysis showing that scene construction could explain the relationship between autobiographical memory and future thinking.

We next performed sensitivity analyses for each of the effects (Table 4 and Figure 2). We first focused on when scene construction was the mediator between autobiographical memory and future thinking (Table 4a, Figures 2a and 2b). As can be seen from the sensitivity values, the indirect effect of scene construction (ρ = .75) was substantially more robust than the direct relationship between autobiographical memory future and thinking (ρ = −.2). On Figure 2 the dashed line represents the average effect, and the plotted line shows what happened to the effect when additional variance is taken into consideration. As can be seen in Figure 2a, the indirect effect only disappeared (i.e. crosses the x axis) when additional variance was very high, compared to the lower variance required for the loss of the direct effect (Figure 2b).

**Figure 2.**
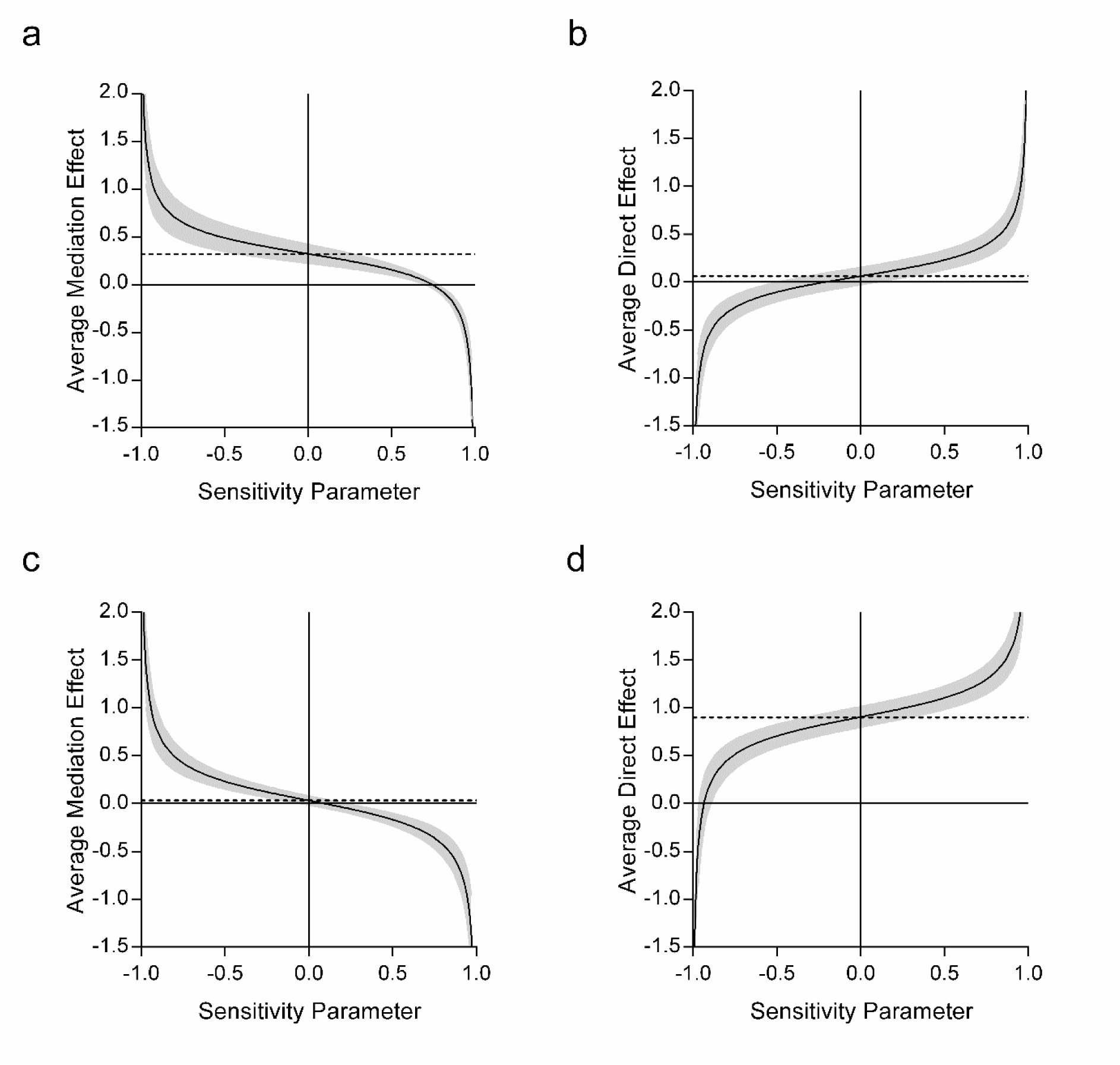
Sensitivity analyses for the indirect and direct effects of the mediation analyses of the Scene component variables, when future thinking was the dependent variable. (a) Sensitivity of the indirect effect of scene construction on the relationship between autobiographical memory and future thinking. (b) Sensitivity of the direct effect between autobiographical memory and future thinking, when scene construction was taken into consideration. (c) Sensitivity of the indirect effect of autobiographical memory on the relationship between scene construction and future thinking. (d) Sensitivity of the direct effect between scene construction and future thinking, when autobiographical memory was taken into consideration. The dashed line shows the average effect when additional variance is assumed to be 0. The plotted line shows the variation in the effect when the additional variance was varied between −1 and 1 (with 95% confidence intervals). The more robust the effect, the greater the variance that was required to reduce the effect to 0 (i.e. to cross the x axis).

A similar, reverse, story was observed when autobiographical memory was used as the mediator between scene construction and future thinking. The autobiographical memory indirect sensitivity rapidly crossed the x axis (ρ = .1, Figure 2c) in comparison to the much higher sensitivity of the direct scene construction to future thinking relationship (ρ = −.95, Figure 2d). Overall, therefore, the effect of scene construction (both as a mediator and directly) was considerably more robust than autobiographical memory, lending additional support to our mediation results. In summary, these first mediation analyses showed that scene construction could explain the relationship between autobiographical memory and future thinking. On the other hand, autobiographical memory could not explain the scene construction future thinking relationship.

Next, we investigated the relationships within the Scene component when future thinking was included as the independent instead of the dependent variable (Table 5, Figure 3). As would be expected, future thinking was associated with both autobiographical memory (Beta = .39, *p* < .001) and scene construction (Beta = .66, *p* < .001). This shows that mediation was possible by both autobiographical memory and scene construction (full regression details are provided in Supplementary Materials, Table S9). However, as before, while the relationship between future thinking and autobiographical memory was fully mediated by scene construction (Table 5a, Figure 3a), the relationship between future thinking and scene construction was only partially mediated by autobiographical memory (Table 5b, Figure 3b). That is, while scene construction could fully explain the relationship between future thinking and autobiographical memory, future thinking was still associated with scene construction even with the additional presence of autobiographical memory.

**Table 5.** Mediation analyses of the Scene component variables when future thinking is the independent variable

|  | Beta (95% CI) | *p* | Sensitivity (*p*) |
| --- | --- | --- | --- |
| a |  |  |  |
| **Future Thinking to Autobiographical Memory, mediated by Scene Construction** |  |  |  |
| Indirect Effect | .25 (.10, .41) | < .001 | .2 |
| Direct Effect | .14 (-.056, .32) | .16 | -.1 |
| Total | .39 (.27, .51) | < .001 | n/a |
| b |  |  |  |
| **Future Thinking to Scene Construction, mediated by Autobiographical Memory** |  |  |  |
| Indirect Effect | .047 (.017, .08) | < .001 | .2 |
| Direct Effect | .62 (.54, .69) | < .001 | -.95 |
| Total | .66 (.59, .73) | < .001 | n/a |

**Figure 3.**
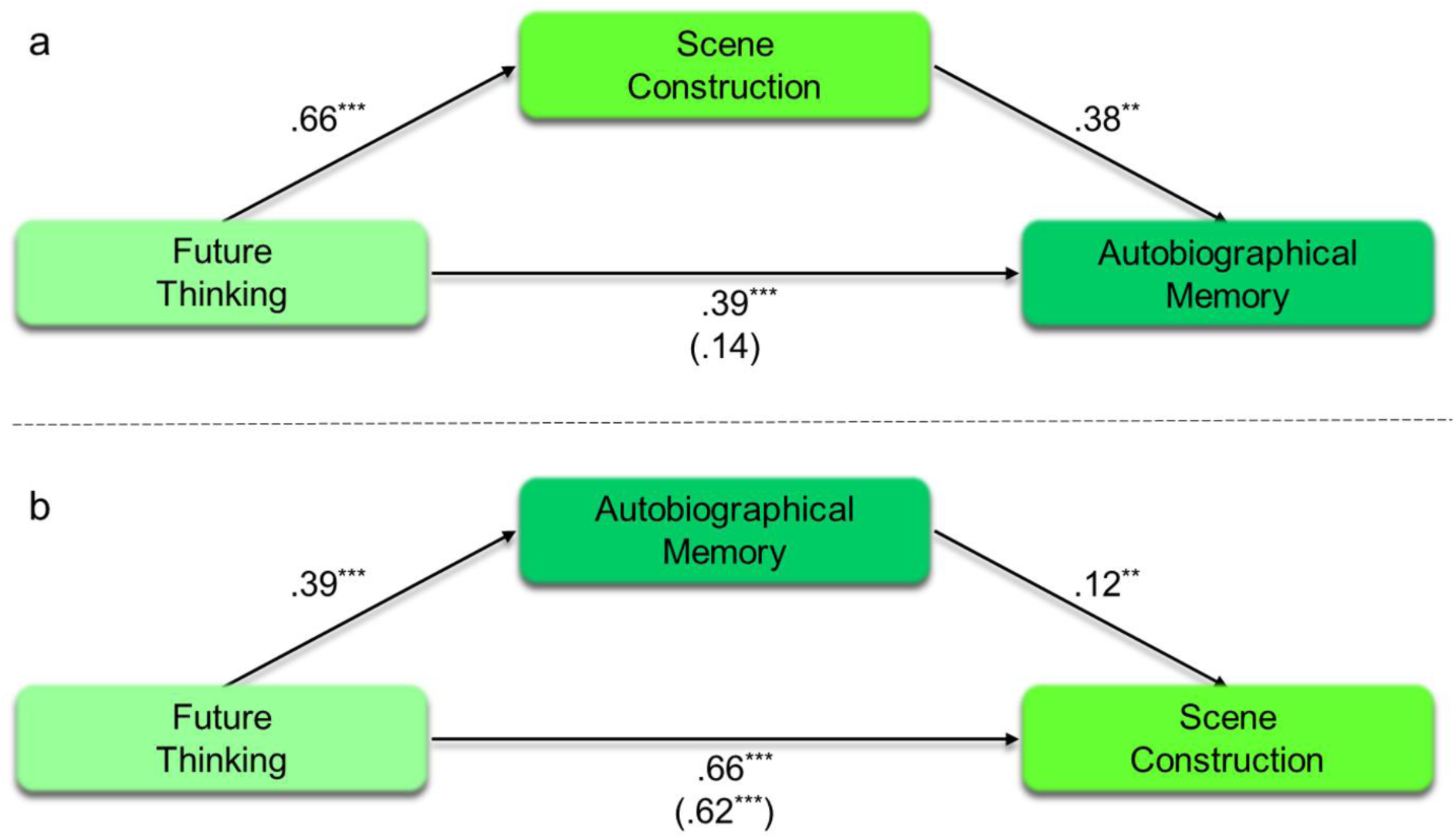
Mediation analyses of the Scene component variables, when future thinking is the independent variable. (a) Future thinking to autobiographical memory, mediated by scene construction. (b) Future thinking to scene construction, mediated by autobiographical memory. The numbers in brackets show the effect of the independent variable on the dependent when the mediation variable was also taken into consideration. ^***^*p* < .001, ^**^*p* <.01.

Looking at the sensitivity analyses, the indirect effect of scene construction on the future thinking-autobiographical memory relationship was small but robust in comparison to the non-significant direct effect of future thinking (ρ = .2 vs. ρ = −.1; Figures 4a and 4b respectively). This supports the mediating role of scene construction on the relationship between future thinking and autobiographical memory. When comparing the sensitivity values for the mediation of autobiographical memory on the future thinking-scene construction relationship, the direct relationship between future thinking and scene construction was much more robust (ρ = −.95, Figure 4d) than the indirect effect of autobiographical memory (ρ = .2, Figure 4c). This highlights that while autobiographical memory may have been contributing something additional to the future thinking-scene construction relationship, it was to a much lesser extent than that of future thinking itself.

**Figure 4.**
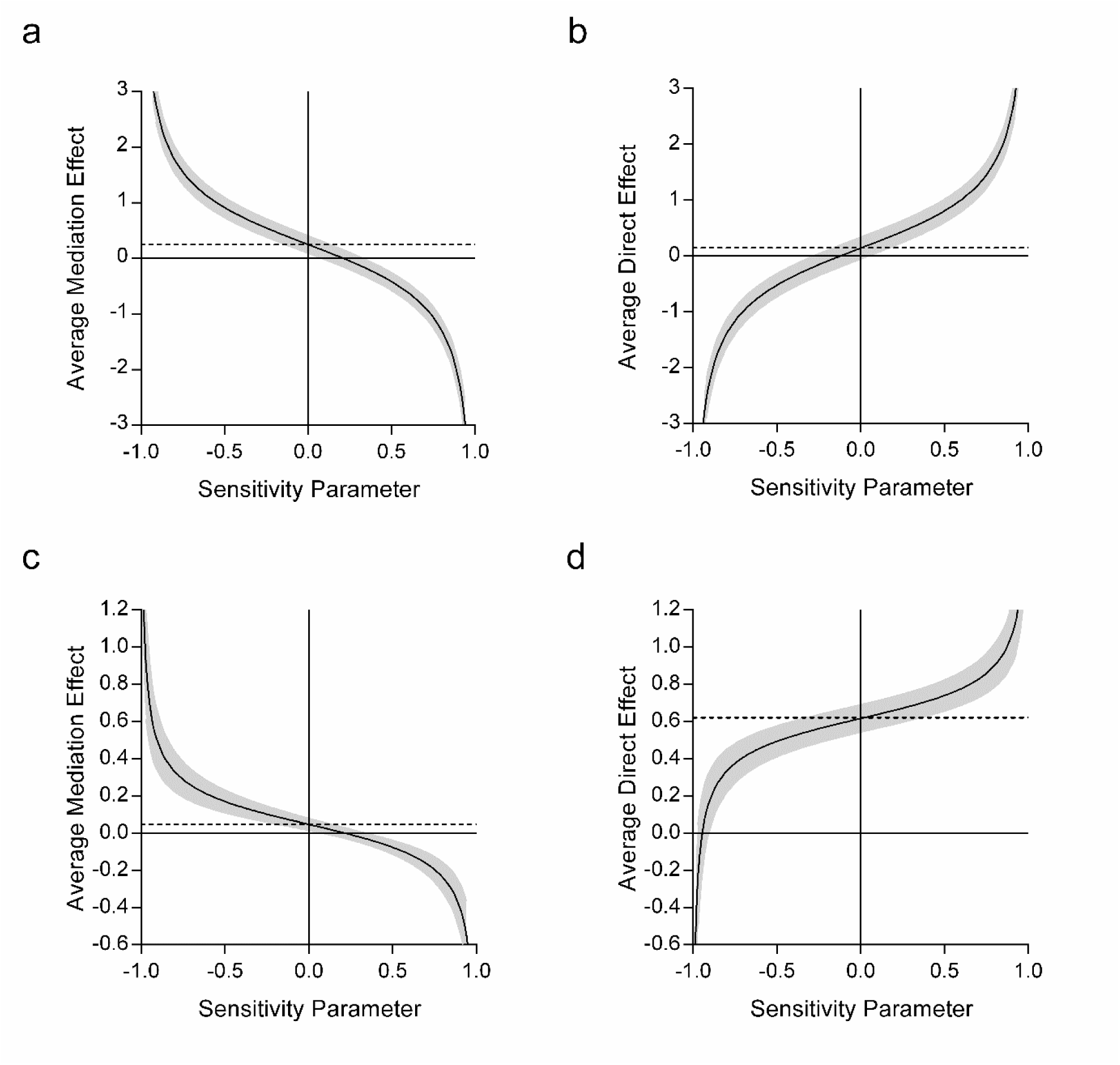
Sensitivity analyses for the indirect and direct effects of the mediation analyses of the Scene component variables when future thinking was the independent variable. (a) Sensitivity of the indirect effect of scene construction on the relationship between future thinking and autobiographical memory. (b) Sensitivity of the direct effect between future thinking and autobiographical memory, when scene construction was taken into consideration. (c) Sensitivity of the indirect effect of autobiographical memory on the relationship between future thinking and scene construction. (d) Sensitivity of the direct effect between future thinking and scene construction, when autobiographical memory was taken into consideration. The dashed line shows the average effect when additional error is assumed to be 0. The plotted line shows the variation in the effect when the additional error is varied between −1 and 1 (with 95% confidence intervals). The more robust the effect, the greater the variance that was required to reduce the effect to 0 (i.e. to cross the x axis).

Finally, for completeness, we also examined whether future thinking mediated the relationship between scene construction and autobiographical memory. While theoretical accounts emphasise either scene construction or autobiographical memory as the underlying cognitive process of future thinking, the reverse remains possible. However, as can be seen in Table 5a and Figure 3a there was no direct effect between future thinking and autobiographical memory with the inclusion of scene construction as the mediator. This, therefore, suggests that future thinking cannot explain the relationship between scene construction and autobiographical memory.

Overall, following a systematic examination of the different configurations of scene construction, autobiographical memory and future thinking, we observed a consistent mediation by scene construction in the various combinations of the relationships between our tasks of primary interest. On the other hand, autobiographical memory seemed to have only limited input. Furthermore, future thinking also failed to explain the relationship between autobiographical memory and scene construction.

In summary, we aimed to assess the underlying psychological process of the Scene component. We predicted that the process of scene construction would best explain the relationships between the three tasks. In line with our prediction, scene construction fully mediated the relationship from autobiographical memory to future thinking and from future thinking to autobiographical memory. Autobiographical memory recall, on the other hand, did not contribute to the relationship from scene construction to future thinking, and only partially mediated the effect of future thinking on scene construction. It seems, therefore, that a key process underpinning the Scene component is indeed related to the mental construction of scene imagery.

### How does the Scene component relate to navigation?

The Scene component only contained three of our tasks of primary interest. The fourth, navigation, aligned instead with the Spatial component. Nevertheless, navigation has long been associated with hippocampal function (Maguire et al., 2000; O’Keefe & Nadel, 1978) and more recently with scene construction (Maguire & Mullally, 2013). Consequently, we also tested whether there was any kind of relationship between the tasks of the Scene component and navigation.

To do this we performed mediation analyses involving scene construction, autobiographical memory, future thinking and navigation. We aimed to establish whether there was an underlying link between the Scene component variables and navigation, predicting that there would be, and that this would be scene construction and not autobiographical memory. As such, we first investigated whether scene construction mediated the relationship between autobiographical memory and navigation. This was then followed by examination of whether autobiographical memory mediated the relationship between scene construction and navigation.

The results of the mediation analyses are shown in Table 6 and Figure 5 (see also Supplementary Materials, Table S10 for the full break down of the regression analyses). Figure 5a shows the relationship between autobiographical memory and navigation with scene construction as the mediator. First, we observed that autobiographical memory was related to both scene construction (Beta = .36, *p* < .001) and navigation (Beta = .99, *p* = .003). This confirmed that mediation by scene construction was possible. Then, importantly, we found that with scene construction included as the mediator, autobiographical memory was no longer related to navigation (Beta = .46, *p* = .2), while scene construction was (Beta = 1.48, *p* < .001). Mediation analysis revealed this to be a significant indirect effect of scene construction, with a non-significant direct effect of autobiographical memory (Table 6a, Figure 5a). This suggested that scene construction fully explained the relationship between autobiographical memory and navigation.

**Table 6.** Mediation analyses of the scene construction, autobiographical memory and navigation relationships

|  | **Beta (95% CI)** | ***p*** | **Sensitivity** |
| --- | --- | --- | --- |
| **a** |  |  |  |
| **Autobiographical Memory to Navigation, mediated by Scene Construction** |  |  |  |
| Indirect Effect | .53 (.21, .90) | < .001 | .25 |
| Direct Effect | .45 (-.25, 1.13) | .21 | -.2 |
| Total | .98 (.32, 1.64) | .003 | n/a |
| **b** |  |  |  |
| **Scene Construction to Navigation, mediated by Autobiographical Memory** |  |  |  |
| Indirect Effect | .23 (-.12, .61) | .20 | .1 |
| Direct Effect | 1.49 (.66, 2.33) | < .001 | -.5 |
| Total | 1.72 (.99, 2.47) | < .001 | n/a |

**Figure 5.**
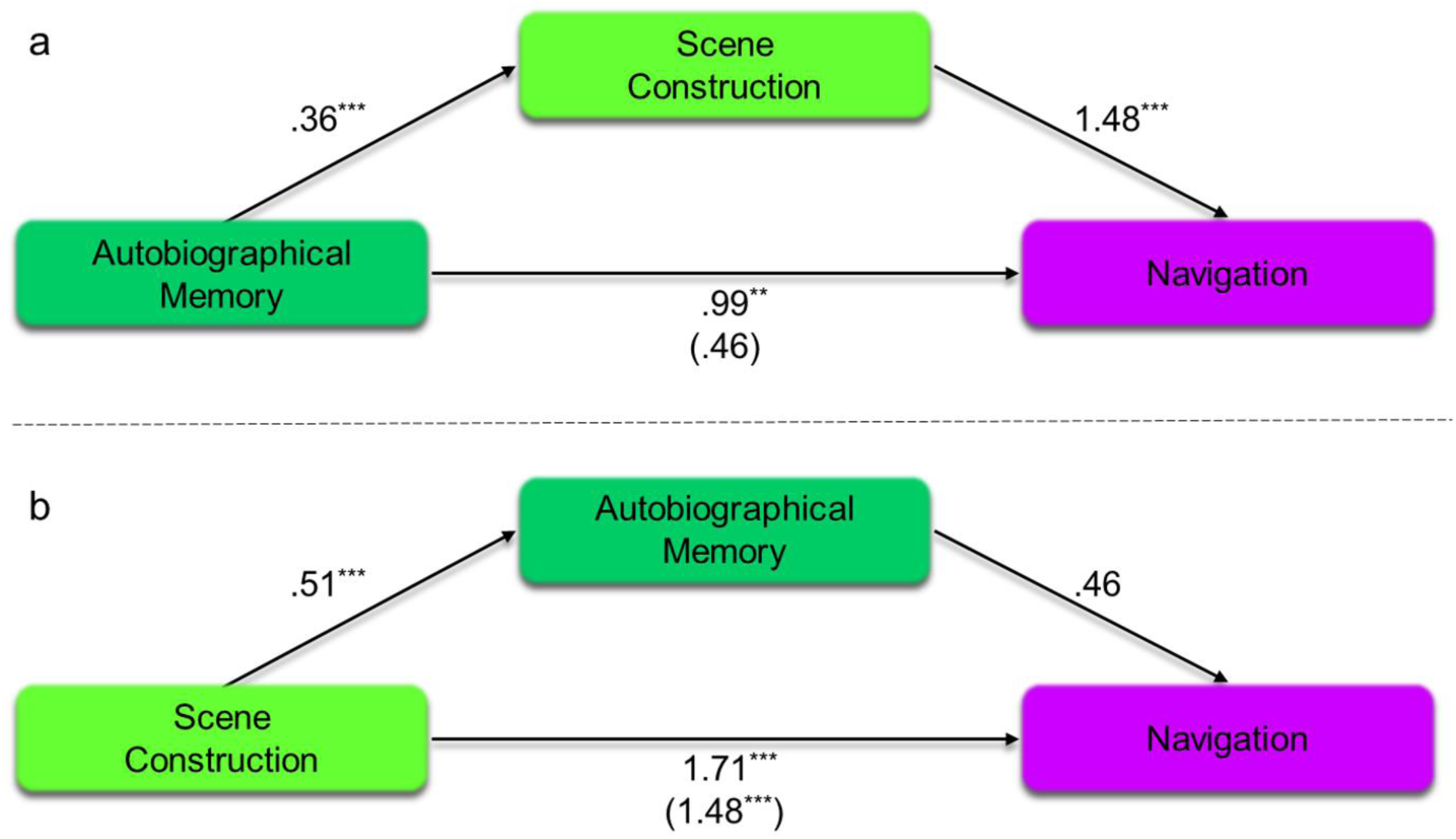
Mediation analyses of the scene construction, autobiographical memory and navigation relationships. (a) Autobiographical memory to navigation, mediated by scene construction. (b) Scene construction to navigation, mediated by autobiographical memory. The numbers in brackets show the effect of the independent variable on the dependent when the mediation variable was also taken into account. ^***^*p* < .001, ^**^*p* < .01.

In contrast, Figure 5b shows the relationship between scene construction and navigation mediated by autobiographical memory. Again we found that scene construction was related to both autobiographical memory (Beta = .51, *p* < .001) and navigation (Beta = 1.71, *p* < .001). As such, mediation by autobiographical memory was possible. However, when autobiographical memory was included as the mediator, no relationship was found between autobiographical memory and navigation (Beta = .46, *p* = .2). Notably, the direct effect between scene construction and navigation remained significant (Beta = 1.48, *p* < .001). Mediation analyses confirmed a significant direct effect in the absence of an indirect effect (Table 6b). This, therefore, suggests that autobiographical memory had no influence on the relationship between scene construction and navigation.

Sensitivity analyses supported both sets of findings, suggesting more robust effects of scene construction than autobiographical memory. This was apparent both when scene construction was mediating the relationship of autobiographical memory and navigation, and for the direct relationship between scene construction and navigation (Table 6, Figure 6). Overall, these results suggest that scene construction may underpin the relationship between autobiographical memory and navigation.

**Figure 6.**
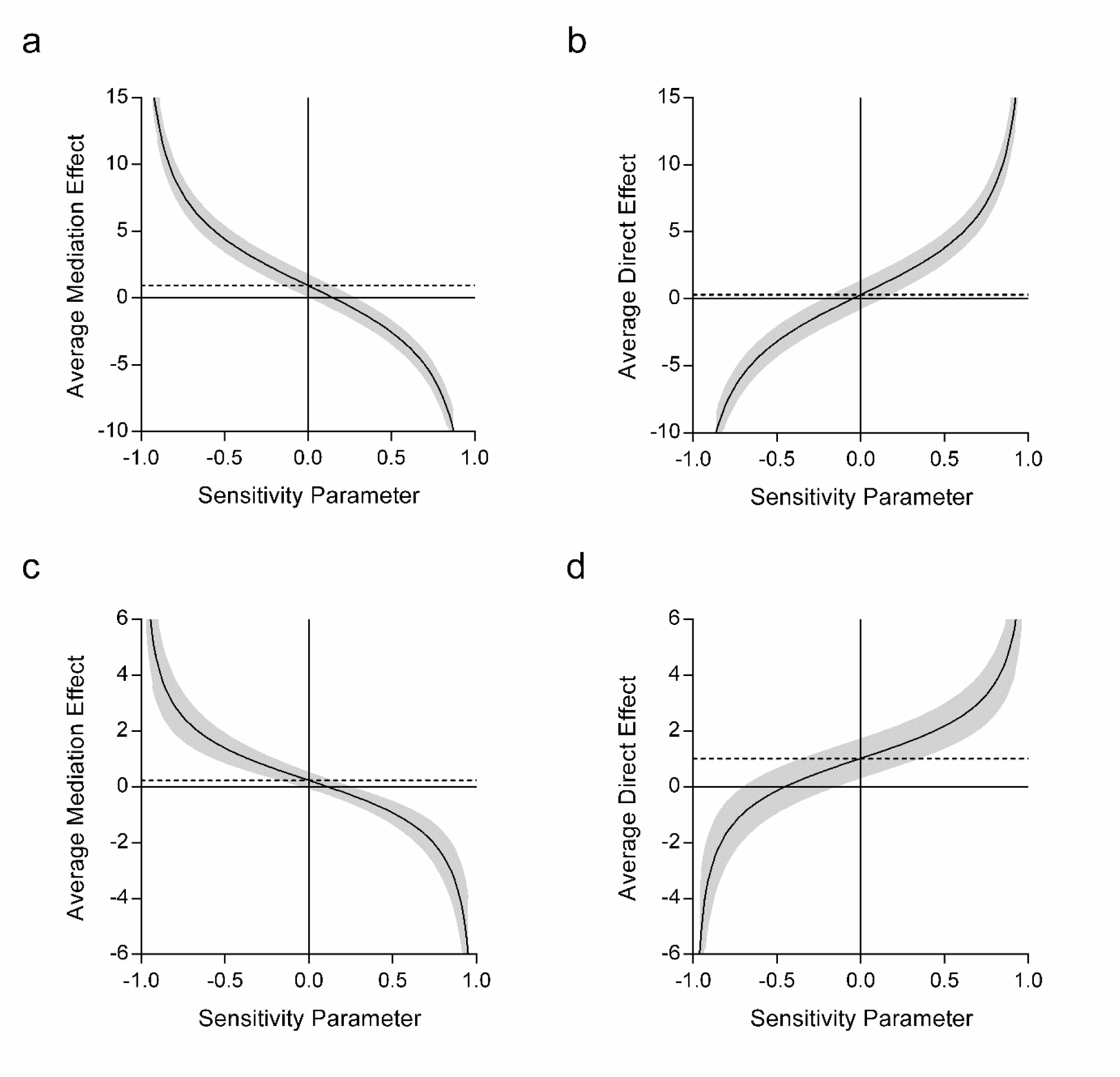
Sensitivity analyses for the indirect and direct effects of the mediation analyses of the scene construction, autobiographical memory and navigation relationships. (a) Sensitivity of the indirect effect of scene construction on the relationship between autobiographical memory and navigation. (b) Sensitivity of the direct effect between autobiographical memory and navigation, when scene construction was taken into consideration. (c) Sensitivity of the indirect effect of autobiographical memory on the relationship between scene construction and navigation. (d) Sensitivity of the direct effect between scene construction and navigation, when autobiographical memory was taken into consideration. The dashed line shows the average effect when additional error is assumed to be 0. The plotted line shows the variation in the effect when the additional error is varied between −1 and 1 (with 95% confidence intervals). The more robust the effect, the greater the variance that was required to reduce the effect to 0 (i.e. to cross the x axis).

We next investigated whether scene construction would also mediate the future thinking to navigation relationship. Once again, we compared this to the mediating ability of autobiographical memory. The results of the mediation analyses are shown in Table 7 and Figure 7 (see Supplementary Materials, Table S11 for the individual regressions). Figure 7a shows the relationship between future thinking and navigation, mediated by scene construction. Future thinking was related to both navigation (Beta = 1.24, *p* < .001) and scene construction (Beta = .66, *p* < .001). This confirmed that mediation by scene construction was possible. With the inclusion of scene construction as the mediator, future thinking was no longer related to navigation (Beta = .29, *p* = .58), while scene construction was (Beta = 1.44, *p* = .022). Mediation analysis identified a significant indirect effect of scene construction, with no direct effect of future thinking (Table 7a). This, therefore, suggests that scene construction fully mediated the relationship between future thinking and navigation, in addition to mediating the autobiographical memory to navigation relationship reported above.

**Table 7.** Mediation analyses of the future thinking to navigation relationship with scene construction or autobiographical memory as the mediating variable

|  | **Beta (95% CI)** | ***p*** | **Sensitivity** |
| --- | --- | --- | --- |
| **a** |  |  |  |
| **Future Thinking to Navigation, mediated by Scene Construction** |  |  |  |
| Indirect Effect | .94 (.11, 1.76) | .025 | .15 |
| Direct Effect | .31 (-.73, 1.33) | .56 | -.05 |
| Total | 1.25 (.60, 1.87) | < .001 | n/a |
| **b** |  |  |  |
| **Future Thinking to Navigation, mediated by Autobiographical Memory** |  |  |  |
| Indirect Effect | .23 (-.034, .54) | .088 | .10 |
| Direct Effect | 1.01 (.29, 1.70) | .0024 | -.45 |
| Total | 1.25 (.60, 1.90) | < .001 | n/a |

**Figure 7.**
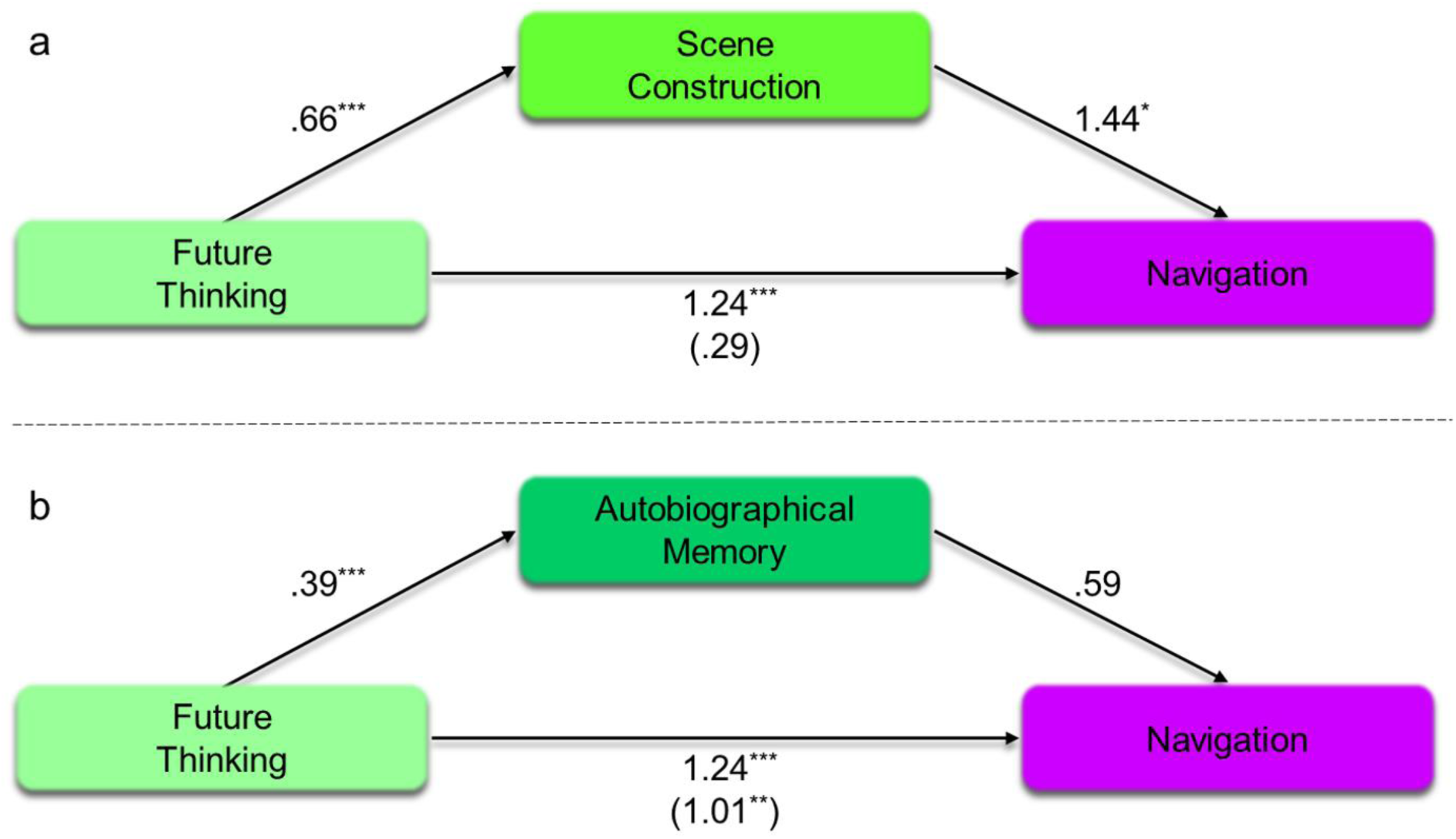
Mediation analyses of the future thinking to navigation relationship with scene construction or autobiographical memory as the mediating variable. (a) Future thinking to navigation, mediated by scene construction. (b) Future thinking to navigation, mediated by autobiographical memory. The numbers in brackets show the effect of the independent variable on the dependent when the mediation variable was also taken into consideration. ^***^*p* < .001, ^**^*p* < .01, ^*^*p* < .05.

On the other hand, Figure 7b shows the relationship between future thinking and navigation, mediated by autobiographical memory. Future thinking was again found to be related to both autobiographical memory (Beta = .39, *p* < .001) and navigation (Beta = 1.24, *p* < .001). This confirmed that mediation by autobiographical memory was possible. However, including autobiographical memory as the mediating variable had limited effect; future thinking remained associated with navigation (Beta = 1.01, *p* = .0045) and there was no relationship between autobiographical memory and navigation (Beta = .59, *p* = .093). Mediation analysis confirmed the absence of an indirect effect of autobiographical memory and the presence of a significant direct effect from future thinking to navigation (Table 7b).

As before, sensitivity analyses were performed to test for the robustness of the effects. These showed, first, a more robust indirect effect of scene construction (ρ = .15, Figure 8a) than the direct relationship between future thinking and navigation (ρ = −.05, Figure 8b). Second, a more robust direct effect of future thinking on navigation (ρ = −.45, Figure 8d) in comparison to the indirect effect of autobiographical memory (ρ = .1, Figure 8c). This supports the mediation analyses.

**Figure 8.**
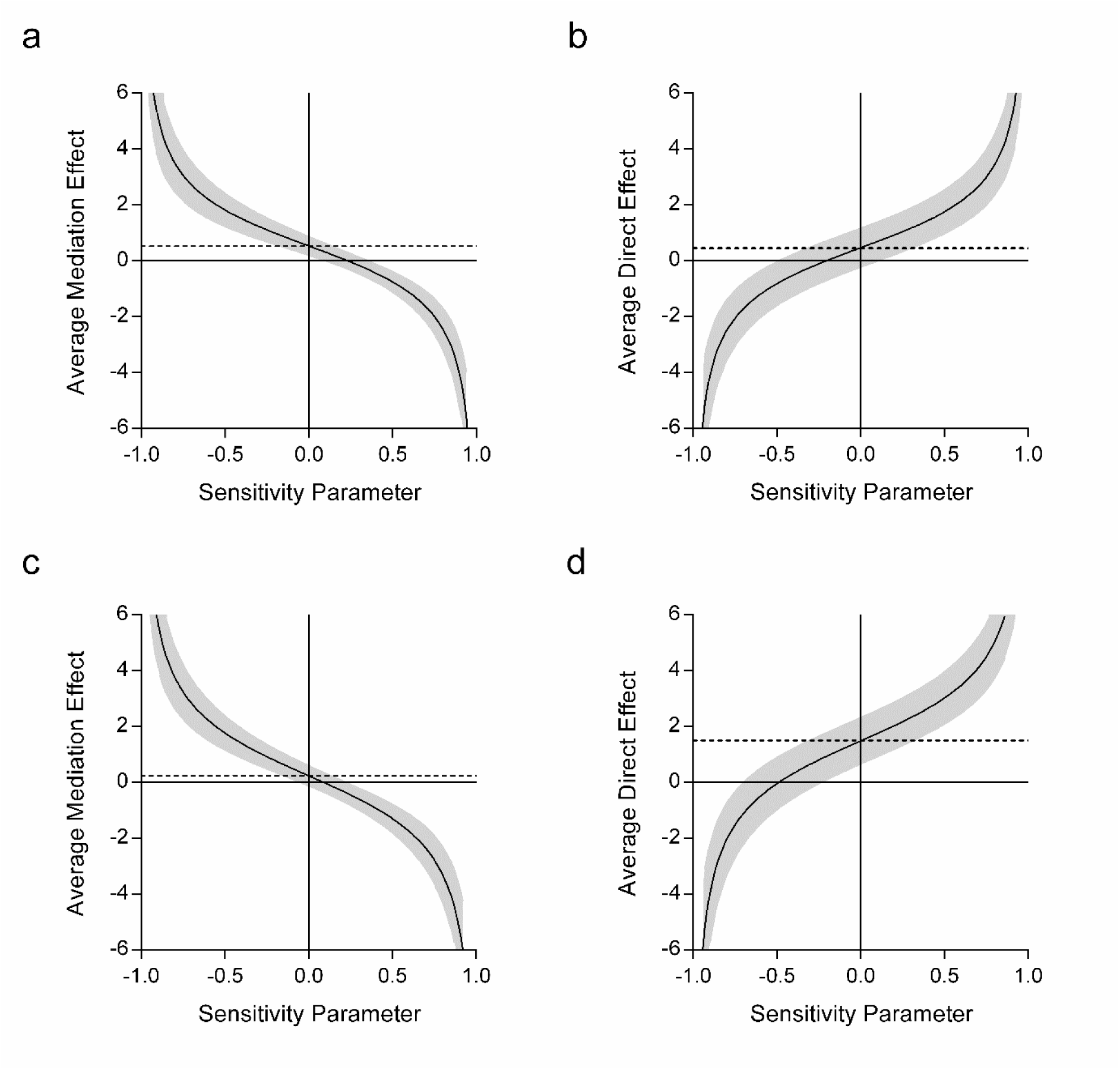
Sensitivity analyses for the indirect and direct effects of the mediation analyses of the future thinking to navigation relationship with scene construction or autobiographical memory as the mediating variable. (a) Sensitivity of the indirect effect of scene construction on the relationship between future thinking and navigation. (b) Sensitivity of the direct effect between future thinking and navigation, when scene construction was taken into consideration. (c) Sensitivity of the indirect effect of autobiographical memory on the relationship between future thinking and navigation. (d) Sensitivity of the direct effect between future thinking and navigation, when autobiographical memory was taken into consideration. The dashed line shows the average effect when additional error is assumed to be 0. The plotted line shows the variation in the effect when the additional error is varied between −1 and 1 (with 95% confidence intervals). The more robust the effect, the greater the variance that was required to reduce the effect to 0 (i.e. to cross the x axis).

We do, however, note that here we have two possible mediators for the future thinking navigation relationship. In addition, the finding of a significant indirect effect of scene construction in comparison to the absence of an indirect effect of autobiographical memory does not necessarily confirm that scene construction is more important than autobiographical memory. We therefore performed an additional analysis with both scene construction and autobiographical memory included as potential mediators on the future thinking navigation relationship at the same time. We found a significant indirect effect of scene construction [Beta = .84 (95% CI: .015, 1.67), *p* = .046] in the absence of an indirect effect of autobiographical memory [Beta = .17 (95% CI: −.10, .45), *p* = .22] and no direct relationship between future thinking and navigation [Beta = .23 (95% CI: −.79, 1.26), *p* = .66]. This, therefore, supports our previous analyses in demonstrating the importance of scene construction, and the absence of the influence of autobiographical memory, in relating the Scene component to navigation.

### Does scene construction retain influence on navigation when spatial processing is taken into account?

The results so far suggest that the process of scene construction may underpin the relationship between our main tasks of primary interest (i.e. scene construction, autobiographical memory, future thinking and navigation). However, it is important to acknowledge that in our initial PCA, navigation loaded on the Spatial component and not the Scene component. This tells us that while scene processing may have some relationship with navigation (as shown by the analyses above), navigation is still closely associated with spatial processing. Consequently, this raises the question of whether scene construction only plays a role in the relationship between the Scene component tasks and navigation in the absence of spatial processing.

To investigate this, we took a similar mediation approach as before, now using the Spatial and Scene components of the PCA. As such, we asked whether the tasks of the Scene component would mediate the relationship between the tasks of the Spatial component and navigation. We did this in two ways. First, we examined the effects of including all three tasks of the Scene component together as a combined mediator variable, to balance the inclusion of the combined Spatial component tasks as the independent variable. Second, we placed scene construction, autobiographical memory and future thinking in turn as separate mediator variables to see if the three tasks of the Scene component had differing mediating effects on the Spatial component to navigation relationship.

Latent variables were used to represent the Scene and Spatial components. The latent variables were comprised of the tasks that loaded singularly onto the respective components. This allowed for assessment of only the pure elements of each component. For the Spatial component this was the ROCF (delayed recall), the Paper Folding test, the object-place association test and the Brixton Spatial Anticipation Test. For the Scene component the tests were scene construction, autobiographical memory and future thinking. To perform a mediation analysis using latent variables, a structural equation modelling (SEM) approach was taken. Aside from the inclusion of latent variables, however, the principles of the analysis remained the same as the mediation analyses reported above. The only exception being that sensitivity analyses can no longer be conducted; judgements are made in SEM on the goodness of model fit.

Figure 9 shows the SEM of the relationship between the Spatial component and navigation, mediated by the Scene component (see also see Supplementary Materials, Table S12 for full details of individual paths). The latent variables (Spatial and Scene PCA components) are shown in circles, the observed variables (the cognitive tasks) in rectangles. The numerical values represent the unstandardised coefficients of the path in question. Overall model fit was good, in line with published recommendations (Hu & Bentler, 1999) [*χ*^2^(18) = 20.70, *p* = .30; CFI = .99; TLI = .99; RMSEA = .026 (90% CI: 0, .068); SRMR = .035]. As would be expected, the ROCF, the Paper Folding test, the object-place association test and the Brixton Spatial Anticipation Test all loaded significantly onto the Spatial latent variable (Beta coefficients respectively of: 3.87, *p* < .001; 2.59, *p* < .001; 1.18, *p* < .001; .66, *p* < .001). Additionally, scene construction, future thinking and autobiographical memory all loaded significantly onto the Scene latent variable (Beta coefficients respectively of: 5.45, *p* < .001; 5.87, *p* < .001; 3.21, *p* < .001). Of key relevance to our question of interest, the Spatial component was associated with the Scene component (Beta = .28, *p* = .002), and both the Spatial and Scene components were associated with navigation (Beta = 22.71, *p* < .001; Beta = 4.87, *p* = .03 respectively). This indicates that the Scene component partially mediated the relationship between the Spatial component and navigation. This is supported by a mediation analysis finding a significant indirect effect of the Scene component (Beta = 1.35 [95% CI: .093, 2.62], *p* = .035). Unsurprisingly, the Spatial component remained associated with navigation even with the introduction of the Scene component.

**Figure 9.**
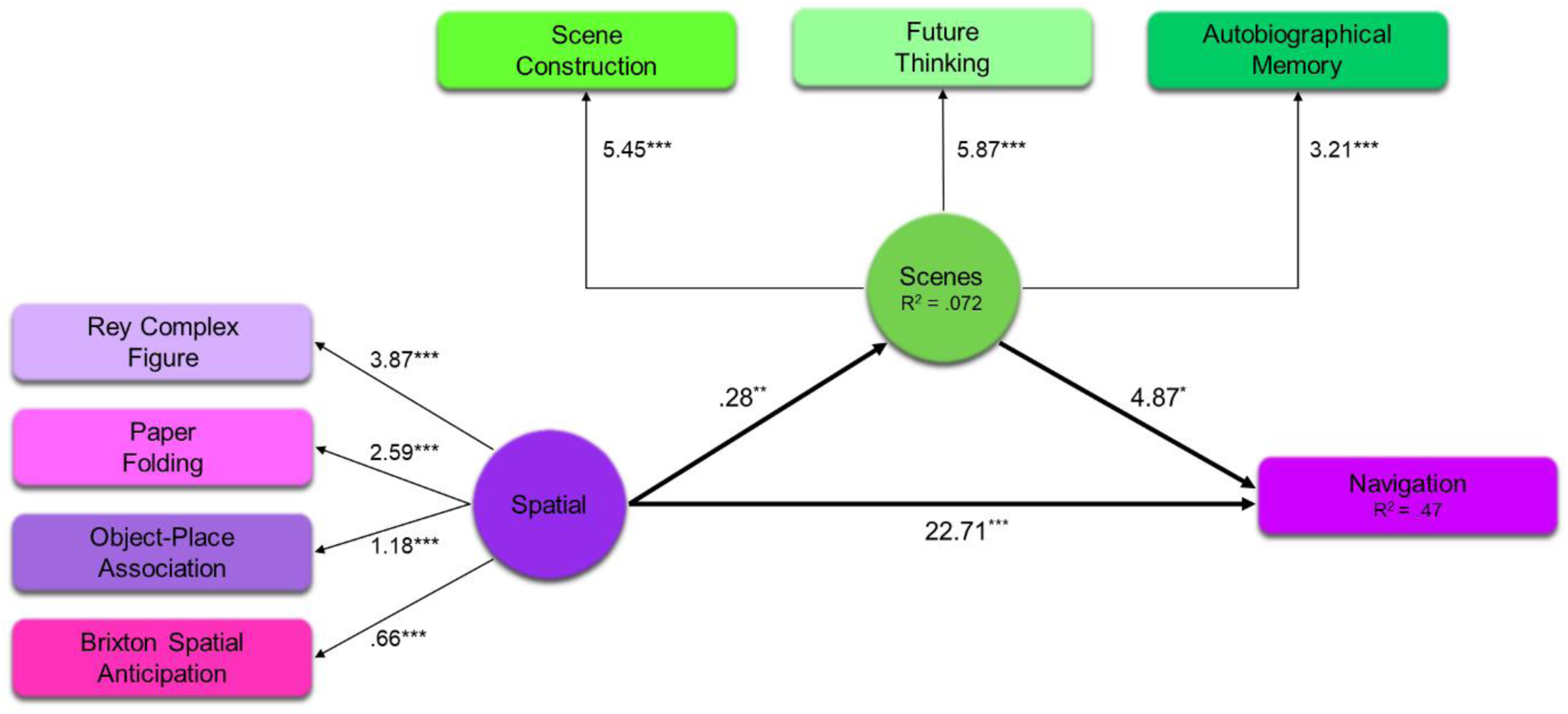
Structural equation model of the Spatial component to navigation relationship mediated by the Scene component. The darker arrows show the main paths of interest, the lighter arrows show the links between the individual observed variables and their related latent variable. The *R*^2^ values represent the proportion of variance explained by the main paths of interest (i.e. the dark arrows). Numerical values linked with a pathway represent unstandardized path coefficients. \**p* < .05, \*\**p* < .01, \*\*\**p* < .001.

Hence, we see a partial mediation by the Scene component in comparison to the full mediations observed earlier. Overall, this suggests that scene processing had an influence on navigation even when the Spatial component was taken into account.

As the Scene component was made up of three variables, we next tested whether the partial mediation by the Scene component on the Spatial component to navigation relationship was specifically due to scene construction or could also be explained by autobiographical memory or future thinking. We therefore repeated the SEM three more times, replacing the Scene component with each individual task in turn. Figure 10 shows the results of the three SEMs using scene construction, autobiographical memory or future thinking as the mediator on the Spatial component to navigation relationship. As before, all models showed acceptable fit [scene construction mediation: *χ*^2^(8) = 14.84, *p* = .062; CFI = .97; TLI = .94; RMSEA = .063 (90% CI: 0, .11); SRMR = .038. Autobiographical memory mediation: *χ*^2^(8) = 15.04, *p* = .058; CFI = .97; TLI = .94; RMSEA = .064 (90% CI: 0, .11); SRMR = .038. Future thinking mediation: *χ*^2^(8) = 15.43, *p* = .051; CFI = .97; TLI = .94; RMSEA = .065 (90% CI: 0, .11); SRMR = .039].

Notably, the patterns of mediation differed in each model. As can be seen in Figure 10a (see also Supplementary Materials, Table S13), when scene construction was used as the mediator, an indirect effect was observed. The Spatial component was associated with scene construction (Beta = 1.48, *p* = .002) and both the Spatial component and scene construction were associated with navigation (Beta = 22.89, *p* < .001; Beta = .79, *p* = .026 respectively). This indicates that, just like the overall Scene component, scene construction partially mediated the relationship between the Spatial component and navigation. This was supported by a mediation analysis finding a significant indirect effect of scene construction (Beta = 1.17 [95% CI: .079, 2.26], *p* = .036).

**Figure 10.**
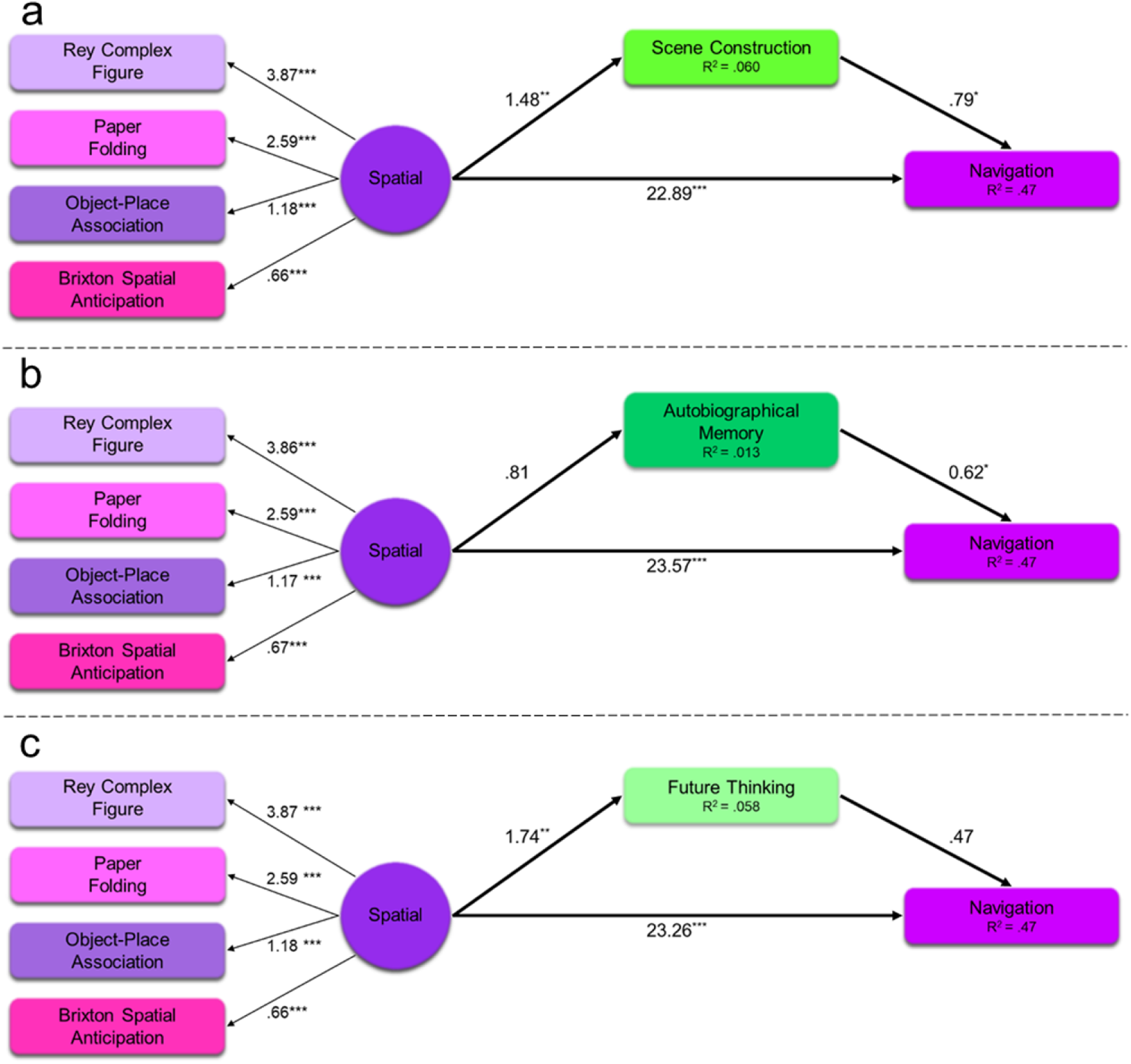
Structural equation models of the Spatial component to navigation relationship mediated by scene construction, autobiographical memory or future thinking. The darker arrows show the main paths of interest, the lighter arrows show the links between the individual observed variables and the latent variable (Spatial). The *R*^2^ values represent the proportion of variance explained by the main paths of interest (i.e. the dark arrows). Numerical values linked with a pathway represent unstandardized path coefficients. \**p* < .05, \*\**p* < .01, \*\*\**p* < .001.

On the other hand, Figure 10b (see also Supplementary Materials, Table S14), shows the effect of using autobiographical memory as the mediator. While the Spatial component continued to be associated with navigation (Beta = 23.57, *p* < .001), the Spatial component was not associated with autobiographical memory (Beta = .81, *p* = .17). As such, while autobiographical memory itself was related to navigation (Beta = .62, *p* = .031), as there was no relationship between the Spatial component and autobiographical memory; these effects were non-mediating. This was supported by the mediation analysis finding no indirect effect of autobiographical memory (Beta = .50 [95% CI: −.23, 1.24], *p* = .18).

Finally, Figure 10c (see also Supplementary Materials, Table S15) shows the indirect effect of future thinking. Here, the Spatial component was associated with both future thinking (Beta = 1.74, *p* = .003) and navigation (Beta = 23.26, *p* < .001). However, there was no relationship between future thinking and navigation when the Spatial component was taken into consideration (Beta = .47, *p* = .12). This suggests that future thinking had no indirect effect on the Spatial component to navigation relationship. This was supported by the mediation analysis (Beta = .81 [95% CI: −.20, 1.83], *p* = .12).

Overall, therefore, we found that scene construction played a role in the relationship between spatial processing and navigation. This is observed by the indirect effects of both the overarching Scene component, and more specifically when just using scene construction. On the other hand, neither autobiographical memory nor future thinking mediated the Spatial component to navigation relationship. To that end, even in the presence of other highly associated spatial tasks, scene construction continued to be a key process involved in navigation.

## Discussion

Autobiographical memory, future thinking, spatial navigation and the imagination of scene imagery are critical cognitive functions that are typically regarded as being related, primarily because they are all hippocampal-dependent. Until now, direct evidence for their interrelatedness has been lacking, as has an understanding of why they might be related. There were four main findings from the current study that spoke to these issues. First, using a PCA, we found that, in the presence of other cognitive tasks, scene construction, autobiographical memory and future thinking all loaded onto the same component, confirming a strong relationship between these variables. Navigation on the other hand, loaded more strongly with spatial tasks. Second, we showed that scene construction fully mediated the relationship between autobiographical memory and future thinking, while autobiographical memory did not mediate between scene construction and future thinking, nor did future thinking mediate between scene construction and autobiographical memory. Third, we found that scene construction fully mediated the relationships between future thinking and autobiographical memory with navigation, while autobiographical memory did not mediate the relationships between future thinking and scene construction with navigation. Finally, we observed a partial mediation by scene construction on the relationship between the spatial tasks and navigation, compared to no mediation of autobiographical memory or future thinking. Overall, our results suggest that scene construction may be a significant cognitive process underlying the relationships between these different functions that are each associated with the hippocampus.

The crucial role of visual imagery is well documented across multiple cognitive domains, including autobiographical memory, future thinking and navigation (Andrews-Hanna, Saxe, & Yarkoni, 2014; Greenberg & Knowlton, 2014; Kraemer et al., 2017). Why might scene imagery in particular be at the heart of these important cognitive functions? One reason is that scene imagery allows us to build models of the world that mirror our moment-by-moment perception. Scenes are also a highly efficient means of packaging information and, as such, are an economical use of cognitive resources (e.g. Konkle, Brady, Alvarez, & Oliva, 2010). Through the construction of a visual scene we can incorporate event details of episodic memories and future events, or route details when navigating, allowing them to be played out in a coherent and naturalistic manner (Maguire & Mullally, 2013; see also Clark & Maguire, 2016).

Revealing the influence of scene construction over autobiographical memory may seem to be in contrast to the decades of work that has strongly associated the hippocampus and autobiographical memory (Cabeza & St. Jacques, 2007; Squire, 1992; Svoboda et al., 2006). We do not deny or diminish this relationship. However, in addition to autobiographical memory, scene construction and thinking about the future have also been associated with the hippocampus (Hassabis, Kumaran, Vann, et al., 2007; Schacter et al., 2012), and there are substantial overlaps in the behavioural correlates of autobiographical memory, scene construction and future thinking (D’Argembeau & Van der Linden, 2004; de Vito et al., 2012; Robin & Moscovitch, 2014). We suggest that our results allow us to start specifying more precisely why these similar, but different, cognitive processes are associated with the hippocampus. In short, our findings point towards scene construction being a common process underlying autobiographical memory and future thinking (Maguire & Mullally, 2013; Zeidman & Maguire, 2016) rather than autobiographical memory being the common component (Addis et al., 2007; Schacter et al., 2012).

It is interesting to note that the PCA loaded navigation with spatial tasks, and not with scene construction, autobiographical memory and future thinking. Navigation also had the smallest effect sizes in terms of the regressions among the primary tasks of interest. Why this is the case will be an interesting topic for future work. For now, we have two speculations. First, imagery comes in multiple forms. A popular distinction is between analytical imagery, reliant upon schematic images, compared to vivid and colourful images of specific scenes and objects (e.g. Kozhevnikov, Kosslyn, & Shephard, 2005). It could be argued that navigation is more like the former, while scene construction, autobiographical memory and future thinking are more similar to the latter. A detailed analysis of the types of imagery being used to perform these tasks may be useful in exploring this further. Second, the distinction between navigation and the other tasks may be because they rely on different hippocampal subregions. Navigation is typically associated with the posterior hippocampus (Maguire et al., 2000), while scene construction, autobiographical memory and future thinking are more often associated with the anterior hippocampus (Dalton & Maguire, 2017; Maguire & Mullally, 2013; Zeidman & Maguire, 2016). Understanding the specialisation of different regions of the hippocampus will also be an important topic for future work.

While the reduced associations with navigation advocate caution in making generalisations from navigation studies to, for example, autobiographical memory, we nevertheless still found that scene construction partially mediated the relationship between the spatial tasks and navigation. Thus, even with navigation being more strongly associated with spatial tasks, the involvement of scene processing remained prominent, whereas, importantly, neither autobiographical memory nor future thinking mediated this relationship.

Here, our main interest was in scene construction, autobiographical memory, future thinking and navigation. As such, the numerous other tasks that were included in the initial PCA are not reported on in detail. However, we make several brief observations in relation to these tests. It is notable that recall and recognition tasks loaded onto separate components, as did episodic and semantic memory tasks. There is still debate in the literature about whether all of these tasks are hippocampal-dependent (Smith et al., 2014; Squire, 1992) or whether only recall and episodic memory tasks require the hippocampus (Eichenbaum, Yonelinas, & Ranganath, 2007). While we do not assess this in detail, our findings are more concordant with this latter perspective.

It is also the case that the recall tasks loaded onto components that were different from the Scene component onto which our primary tasks of interest loaded. If the hippocampus is involved in supporting memory recall tasks and also scene construction, autobiographical memory and future thinking, why did they all not cluster onto one factor? The data suggest that the standardised tests in particular clustered according to the modality in which a test was presented. That is, all the verbal recall tasks loaded together, and the visual recall tasks loaded on the spatial component. This does not mean that these tasks are unrelated to our primary tasks of interest, but rather that modality exerted a significant influence.

One question that is often asked of the scene construction theory, is why does hippocampal damage result in verbal memory deficits, for example, in word paired associates tasks, if visuospatial scenes are of particular relevance for hippocampal function? We have previously suggested that some verbal tasks may in fact engage scene imagery (e.g. imagining the two objects in a word pair together in a scene; Clark & Maguire, 2016; Maguire & Mullally, 2013), and that this could explain their dependence on the hippocampus. Recent work using functional neuroimaging lends credence to this idea by finding that high imagery concrete word pairs evoked hippocampal activity due to the use of scene imagery, while low imagery abstract word pairs did not (Clark et al., 2018). Another way to test this in the future would be to interrogate the explicit strategies that people use to perform different verbal recall tasks. This would enable us to ascertain if scene imagery is involved more generally in verbal tasks, and indeed whether the use of such imagery confers a performance advantage.

We note that the Scene component of the PCA contained tasks that were scored from open ended verbal descriptions. As such, verbal task demands - be that narrative style, verbal ability and so forth - or similarities in scoring across the tasks could be candidate processes linking scene construction, autobiographical memory and future thinking. However, if this was the case, we would have expected a different pattern of results to emerge. First, autobiographical memory external details should have loaded onto the Scene component, and it did not. Second, the loading of the scene description task should have been stronger, more in line with the loadings of scene construction, future thinking and autobiographical memory, but it was not. Finally, future thinking should have mediated the relationship between autobiographical memory and scene construction and the relationship between the Spatial component and navigation, and yet it did not. Instead, we observed that external details loaded onto the Semantic Memory component, that the scene description task loaded most strongly on the Perception component and that there was only a mediating effect of scene construction.

In addition, to further examine the potential involvement of verbal processing, we also ran a series of control mediation analyses looking at the effects of the Verbal Memory component (as a proxy for verbal ability) on the tasks of the Scene component (see Supplementary Materials and Figure S2). We found that the influence of the Verbal Memory component was either fully or partially mediated in all the models. This suggests that the relationships between scene construction, autobiographical memory and future thinking we reported above cannot simply be explained by verbal ability.

A related potential criticism is that we tested the influence of memory on the relationship between scene construction, autobiographical memory and future thinking using a memory task that relied on verbal output. To address this issue, we also examined the relationships between scene construction, autobiographical memory and future thinking with a nonverbal memory task (the delayed recall of the ROCF) as the mediator, and found no influence on any of the relationships (see Supplementary Materials, Tables S15–16). This lends further support to the idea that scene imagery rather than memory may be a key feature underlying the relationships between scene construction, autobiographical memory and future thinking.

Finally, we also observed the surprising finding in the PCA analysis that the Brixton Spatial Anticipation Test, Matrix Reasoning, and the Symbol Span test loaded most strongly on the Spatial component. This was unexpected because these tasks are typically thought to tax executive functioning and general intellectual ability (e.g. Wechsler, 2008, 2009). Studies using these standardised tasks should perhaps bear this in mind, as our data suggest that individual differences in spatial processing could affect performance on these tasks.

Here we have alluded to the function of the hippocampus without measuring the hippocampus itself. We feel confident in doing so because of the many previous findings associating the hippocampus with scene construction, autobiographical memory, future thinking and navigation. Moreover, the issue of central interest here – to understand the cognitive processes involved in these tasks – is not reliant upon direct hippocampal measurement. However, an important next step will undoubtedly be to directly relate the process of scene construction with structural and functional measurements of the hippocampus.

We also acknowledge that scene construction, autobiographical memory, future thinking and navigation have each been associated with brain regions outside of the hippocampus including (but not limited to) parahippocampal, retrosplenial, posterior cingulate, parietal and medial prefrontal cortices (e.g. Hassabis, Kumaran, & Maguire, 2007; Schacter et al., 2012; Stawarczyk & D’Argembeau, 2015). For example, lesions to the parietal cortex impair the subjective experience associated with autobiographical memory (Ciaramelli et al., 2017; Simons, Peers, Mazuz, Berryhill, & Olson, 2010) and posterior parietal damage has been linked to a reduction in scene construction ability (Ramanan et al., 2018). A variety of extra-hippocampal brain regions have been implicated in supporting navigation, and a reliance upon non-hippocampal regions for navigation has been suggested to increase with age (Moffat, Elkins, & Resnick, 2006; Zhong & Moffat, 2018). An important future step will, therefore, be to understand the interactions, both structural and functional, between the hippocampus and these other regions, and their relationships with individual differences in task performance.

We also note that it is unlikely the hippocampus supports only one fundamental process. A number of studies have associated different hippocampal subregions with distinct cognitive processes (e.g. Dalton, Zeidman, McCormick, & Maguire, 2018; Dimsdale-Zucker, Ritchey, Ekstrom, Yonelinas, & Ranganath, 2018; Hodgetts et al., 2017; Zeidman, Lutti, & Maguire, 2015). The results from the current study suggest that the construction of scene imagery seems to plays an influential role in autobiographical memory, future thinking and, to some extent, navigation, and consequently, scene construction may be at least one process performed by the hippocampus.

In conclusion, we are not alone in suggesting that the hippocampus is more than just a memory device (O’Keefe & Nadel, 1978; Shohamy & Turk-Browne, 2013; Tulving, 2002; Verfaellie & Keane, 2017). However, here, a large sample of participants, numerous cognitive tests and a wide variance in performance enabled us to provide novel evidence regarding the interrelations between tasks that have hitherto not been systematically examined. We found that the construction of scene imagery plays a particularly prominent role in several hippocampal-dependent tasks. This finding lays the groundwork for future studies that should directly examine the strategies and types of imagery people use to perform such tasks, and how this is realised by the hippocampus and its specific subregions.

### Context

The current study is part of a body of work investigating the relationships between a diversity of tasks that have individually been associated with a brain structure called the hippocampus, located deep in the brain’s temporal lobes. There is debate about the hippocampus’ contribution to these tasks which, on face value, appear to be distinct. Here, we highlight the importance of visual scene imagery for three tasks typically associated with the hippocampus – autobiographical memory, future thinking and spatial navigation. Future work aims to build on these findings by conducting a detailed analysis of the explicit strategies deployed by participants to perform these tasks and whether scene imagery plays a role, and by examining how variations in task performance may be related to structural and functional measurements of the brain, including the hippocampus.

## Clark et al. Supplementary Materials

### Supplementary Methods

**Table S1.** Double scoring of the scene construction test

|  | Rating |  |  |  |  |
| --- | --- | --- | --- | --- | --- |
|  | Spatial<br>References | Entities<br>present | Sensory<br>Descriptions | Thoughts/<br>Emotions/Actions | Quality<br>ratings |
| **For each individual scene** |  |  |  |  |  |
| n = 308 | .90 | .96 | .94 | .90 | .90 |
| **For each individual participant (i.e. score is averaged across the seven scenes)** |  |  |  |  |  |
| n = 44 | .91 | .99 | .97 | .91 | .93 |

*Note.* Inter-class correlation coefficients from a two way random effect model looking for absolute agreement for each content score and for the quality ratings. Four experimenters scored the whole data set (n = 217 participants, 1519 individual scenes) with double scoring performed on 20% of the data (n = 44 participants, 308 scenes) proportionally for each original experimenter.

**Table S2.**
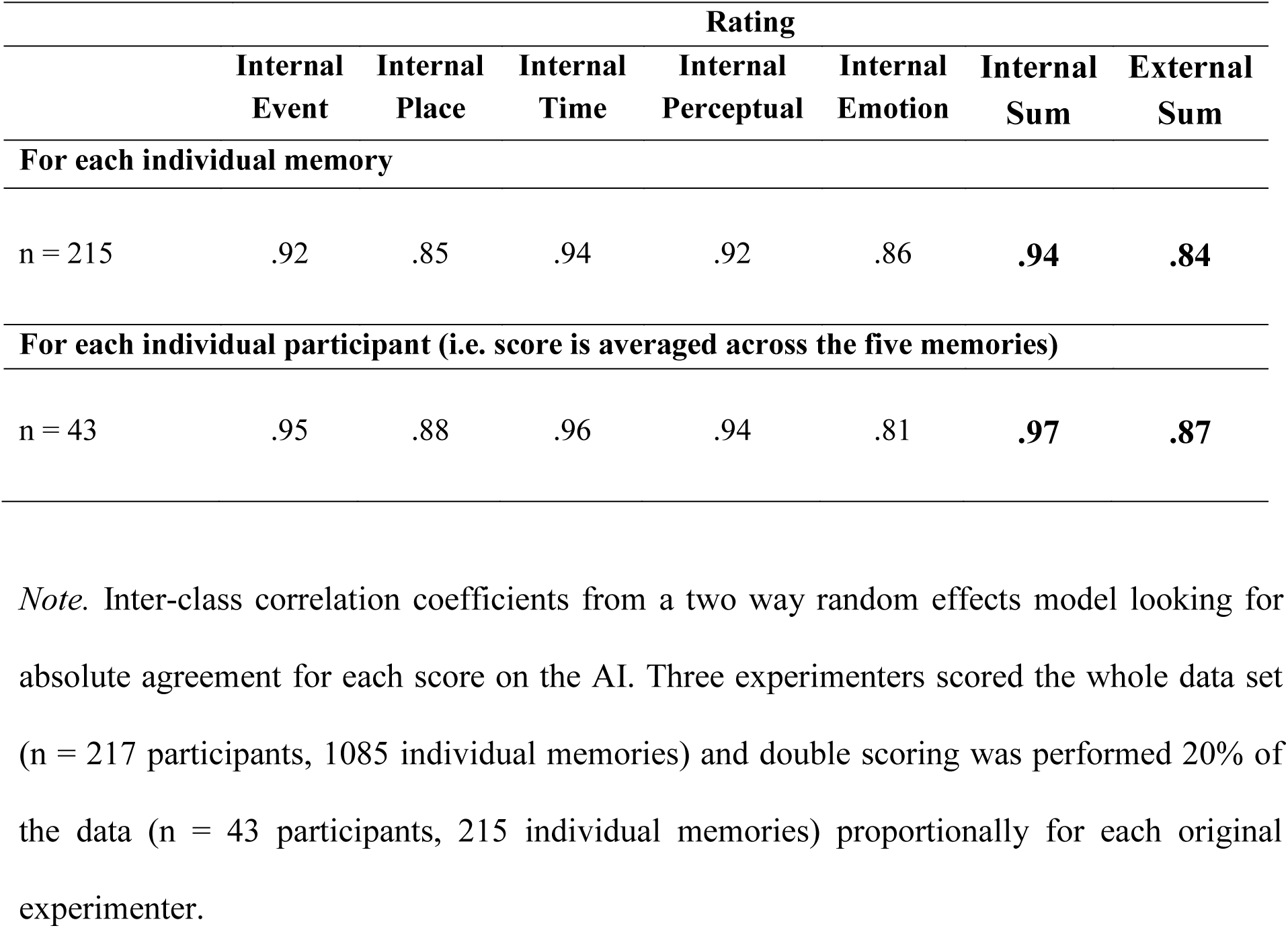
Double scoring of the Autobiographical Interview

|  | Rating |  |  |  |  |  |  |
| --- | --- | --- | --- | --- | --- | --- | --- |
|  | Internal<br>Event | Internal<br>Place | Internal<br>Time | Internal<br>Perceptual | Internal<br>Emotion | Internal<br>Sum | External<br>Sum |
| **For each individual memory** |  |  |  |  |  |  |  |
| n = 215 | .92 | .85 | .94 | .92 | .86 | **.94** | **.84** |
| **For each individual participant (i.e. score is averaged across the five memories)** |  |  |  |  |  |  |  |
| n = 43 | .95 | .88 | .96 | .94 | .81 | **.97** | **.87** |

*Note.* Inter-class correlation coefficients from a two way random effects model looking for absolute agreement for each score on the AI. Three experimenters scored the whole data set (n = 217 participants, 1085 individual memories) and double scoring was performed 20% of the data (n = 43 participants, 215 individual memories) proportionally for each original experimenter.

**Table S3.** Double scoring of the future thinking test

|  | Rating |  |  |  |  |
| --- | --- | --- | --- | --- | --- |
|  | Spatial<br>References | Entities<br>present | Sensory<br>Descriptions | Thoughts/<br>Emotions/Actions | Quality<br>ratings |
| **For each individual scene** |  |  |  |  |  |
| n = 132 | .90 | .94 | .93 | .88 | .90 |
| **For each individual participant (i.e. score is averaged across the three future scenes)** |  |  |  |  |  |
| n = 44 | .94 | .95 | .96 | .88 | .92 |

*Note.* Inter-class correlation coefficients from a two way random effects model looking for absolute agreement for each content score and for the quality ratings. Four experimenters scored the whole data set (n = 217 participants, 651 individual future scenes) with double scoring performed on 20% of the data (n = 44 participants, 132 future scenes) proportionally for each original experimenter.

**Table S4.** Double scoring of the navigation sketch maps

|  | **Rating** |  |  |  |  |  |
| --- | --- | --- | --- | --- | --- | --- |
|  | **Road**<br>**Segments** | **Road**<br>**Junctions** | **Number of**<br>**Landmarks** | **Landmark**<br>**Placement** | **Map**<br>**Orientation** | **Map**<br>**Categorisation** |
| n = 42 | .95 | .96 | .97 | .96 | .96 | .89 |

*Note.* Inter-class correlation coefficients from a two way random effects model looking for absolute agreement for each score on the navigation sketch maps. Three experimenters scored the whole data set (n = 217) and double scoring was performed on 20% of the data (n = 42 participants) proportionally for each original experimenter.

**Table S5.** Double scoring of the scene description task

|  | **Rating** |  |  |  |
| --- | --- | --- | --- | --- |
|  | **Spatial**<br>**References** | **Entities**<br>**present** | **Sensory**<br>**Descriptions** | **Thoughts/**<br>**Emotions/Actions** |
| n = 43 | .88 | .91 | .93 | .85 |

*Note.* Inter-class correlation coefficients from a two way random effects model looking for absolute agreement for each content score. Three experimenters scored the whole data set (n = 217) with double scoring performed on 20% of the data (n = 43) proportionally for each original experimenter.

### Supplementary Results

#### How are the tasks interrelated? Additional results of the PCA

The selection of the number of components in the solution of the PCA was performed using two methodologies. First, as detailed in the main body of the text, a minimum Eigen value set to one suggested a solution of seven components. However, as some of these components contained only two or three variables, we sought to validate the seven factor solution via a second methodology for component selection. This second approach used the “elbow” of the Scree plot of the Eigen values of each component. The Scree plot elbows detail where the explanatory power of the components begins to reduce, and thus become less useful for understanding the relationships in the data. However, it is not always clear where the elbow of the Scree plot is, and so potential solutions also have to be checked for interpretability. Choosing too few components may miss valid relationships in the data, while choosing too many can lead to the inclusion of components that account for only trivial amounts of variance and reduce the interpretability of more interesting components. As can be seen in Figure S1, the elbow of the Scree plot for the PCA could be placed in three locations, suggesting solutions of two, four or seven components. To identify the best solution, we looked at the interpretability of the three solutions.

When limiting the solution to two components, only 32% of the variance was explained. More importantly, two tasks – the Dead or Alive task and the boundary extension task – did not load on either component (highest loadings of .19 and .26 respectively). A two component solution was therefore deemed unsatisfactory and discounted.

When limiting to four components, the total variance explained remained low at only 45%. Importantly, once again, neither the Dead or Alive task nor the boundary extension task loaded onto any of the components (highest loadings of .26 and .27 respectively). A four component solution was therefore also deemed unsatisfactory and discounted.

However, as detailed in the main text, a solution of seven components not only explained 59% of the total variance but, crucially, loaded both the Dead or Alive and boundary extension tasks onto specific components of the PCA (loadings of .62 and .84 respectively). As such, both the minimum Eigen value methodology and the Scree plot methodology suggested the optimal solution of the PCA was that of seven components.

**Figure S1.**
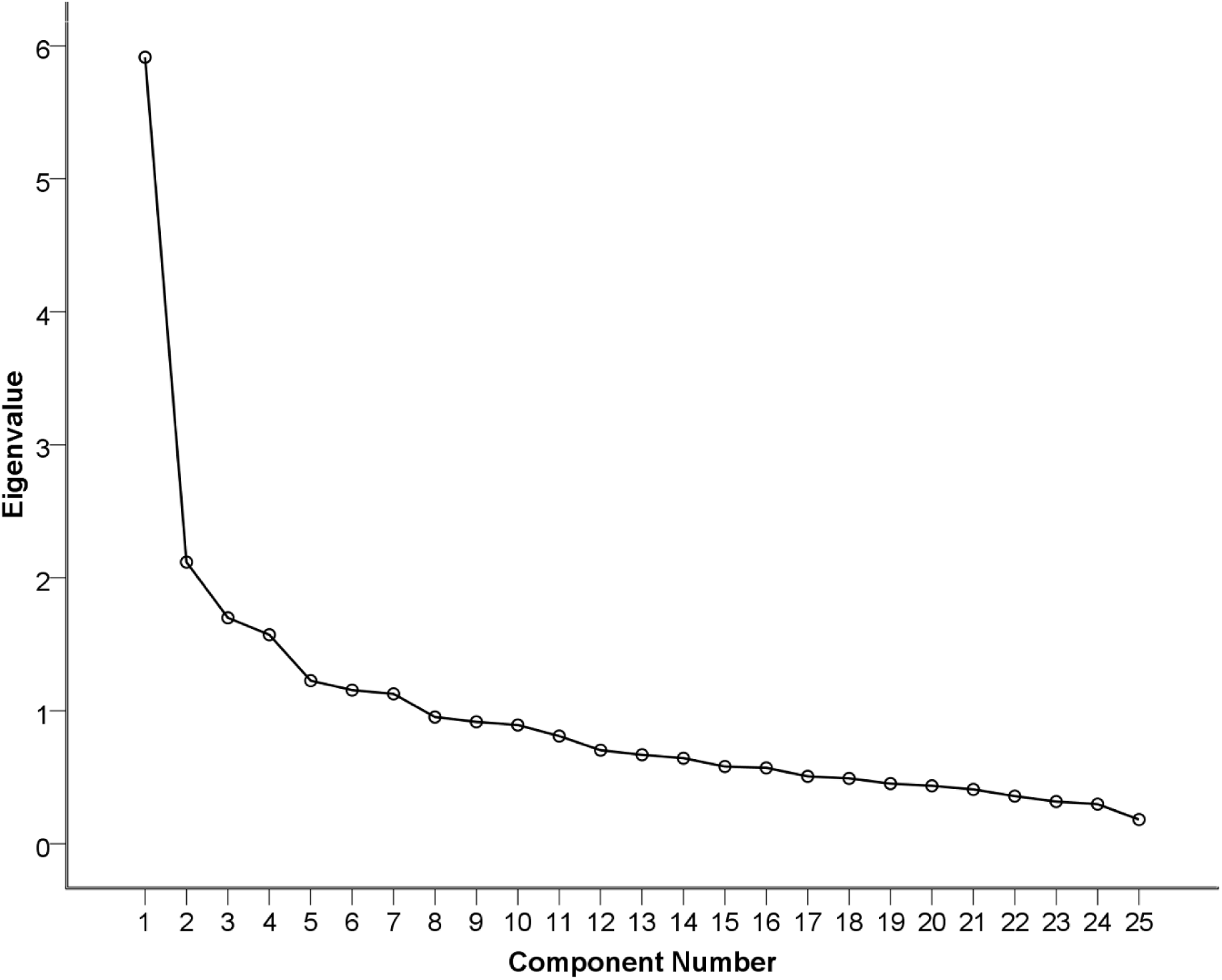
Scree plot of the Principal Component Analysis showing the Eigen values of the proposed components.

**Table S6.** Full details of the Principal Component Analysis with varimax rotation of the cognitive tasks. Task order is for display purposes only

| Cognitive Task | Spatial | Verbal | IQ/Executive Function | Scenes | Recognition Memory | Semantic Memory | Perception |
| --- | --- | --- | --- | --- | --- | --- | --- |
| Rey-Osterrieth Complex Figure delayed recall | .72 | .26 | .022 | .075 | .023 | -.014 | .18 |
| Paper Folding Test | .72 | .18 | .20 | -.014 | -.047 | .13 | -.073 |
| Navigation | .66 | .11 | .17 | .20 | .17 | -.054 | .10 |
| Object-Place Association Test | .65 | .16 | -.096 | .11 | .11 | -.22 | -.072 |
| Brixton Spatial Anticipation Test | .42 | -.13 | .34 | -.16 | .20 | .18 | -.25 |
| Warrington Recognition Memory Test for Scenes | .54 | .025 | .12 | .13 | .41 | .15 | -.045 |
| Matrix Reasoning | .51 | .069 | .49 | .047 | -.011 | .11 | -.072 |
| Symbol Span | .46 | .29 | .46 | .045 | .065 | .031 | -.005 |
| Rey Auditory Verbal Learning Test delayed recall | .21 | .74 | .063 | .064 | -.002 | .066 | .098 |
| Concrete Verbal Paired Associates delayed recall | .20 | .67 | .20 | .039 | .27 | .25 | -.008 |
| Logical Memory delayed recall | .18 | .66 | .048 | .15 | .005 | -.16 | .032 |
| Verbal Paired Associates delayed recall | .13 | .62 | .17 | .085 | .33 | .061 | -.19 |
| Abstract Verbal Paired Associates delayed recall | .033 | .61 | .46 | -.031 | .13 | .035 | -.043 |
| Digit Span | .16 | .17 | .74 | .046 | -.21 | -.17 | -.098 |
| Full Scale Intelligence Quotient | .024 | .20 | .68 | .075 | .26 | .20 | .20 |
| F-A-S Verbal Fluency | .13 | .11 | .62 | .27 | .062 | .027 | .053 |
| Scene Construction Experiential Index | .16 | .028 | .12 | .87 | .072 | .12 | -.065 |
| Future Thinking Experiential Index | .15 | .006 | .16 | .85 | .091 | .084 | -.023 |
| Autobiographical Memory Internal Details | .024 | .24 | -.001 | .62 | -.010 | .13 | .16 |
| Scene Description | -.094 | .036 | .20 | .37 | .19 | -.24 | .50 |
| Warrington Recognition Memory Test for Words | .16 | .18 | -.084 | .095 | .67 | -.18 | .008 |
| Warrington Recognition Memory Test for Faces | .080 | .15 | .11 | .028 | .79 | .089 | .061 |
| Autobiographical Memory External Details | .060 | -.089 | -.051 | .27 | -.060 | .69 | .17 |
| Dead or Alive Task | -.028 | .15 | .087 | .051 | .028 | .62 | -.067 |
| Boundary Extension | .059 | -.036 | -.30 | -.038 | -.015 | .14 | .84 |
| **Variance explained (Total = 59.24%)** | **12.44** | **10.64** | **9.85** | **9.23** | **6.81** | **5.28** | **5.01** |

**Table S7.** Full details of the Principal Component Analysis with FSIQ removed. Task order is for display purposes only

| Cognitive Task | Spatial | Verbal | Scenes | IQ/Executive Function | Recognition Memory | Semantic Memory | Perception |
| --- | --- | --- | --- | --- | --- | --- | --- |
| Rey-Osterrieth Complex Figure delayed recall | .71 | .25 | .062 | .080 | .027 | 0.002 | .19 |
| Paper Folding Test | .70 | .17 | -.036 | .27 | -.038 | .16 | -.052 |
| Navigation | .68 | .13 | .21 | .15 | .13 | -.060 | .071 |
| Object-Place Association Test | .67 | .15 | .10 | -.062 | .086 | -.21 | -.093 |
| Brixton Spatial Anticipation Test | .40 | -.12 | -.16 | .35 | .19 | .21 | -.25 |
| Warrington Recognition Memory Test for Scenes | .56 | .034 | .15 | .097 | .38 | .14 | -.065 |
| Matrix Reasoning | .43 | .065 | .021 | .60 | .050 | .15 | -.005 |
| Symbol Span | .43 | .30 | .056 | .50 | .075 | .037 | .007 |
| Rey Auditory Verbal Learning Test delayed recall | .21 | .73 | .067 | .060 | .011 | .056 | .12 |
| Concrete Verbal Paired Associates delayed recall | .18 | .67 | .034 | .19 | .29 | .26 | .010 |
| Logical Memory delayed recall | .23 | .68 | .16 | -.027 | -.036 | -.16 | -.037 |
| Verbal Paired Associates delayed recall | .11 | .61 | .074 | .19 | .36 | .086 | -.17 |
| Abstract Verbal Paired Associates delayed recall | .009 | .63 | -.013 | .43 | .16 | .027 | -.022 |
| Digit Span | .11 | .20 | .064 | .76 | -.17 | -.16 | -.073 |
| F-A-S Verbal Fluency | .06 | .12 | .28 | .67 | .12 | .040 | 0.12 |
| Scene Construction Experiential Index | .17 | .030 | .86 | .10 | .059 | .13 | -.086 |
| Future Thinking Experiential Index | .15 | .009 | .85 | .15 | .087 | .099 | -.038 |
| Autobiographical Memory Internal Details | .025 | .24 | .62 | -.009 | -.007 | .11 | .18 |
| Scene Description | -.053 | .023 | .42 | .15 | .23 | -.18 | .46 |
| Warrington Recognition Memory Test for Words | .17 | .15 | .080 | -.076 | .69 | -.16 | .010 |
| Warrington Recognition Memory Test for Faces | .085 | .15 | .048 | .073 | .79 | .077 | .073 |
| Autobiographical Memory External Details | .065 | -.090 | .27 | -.075 | -.071 | .69 | .18 |
| Dead or Alive Task | -.038 | .16 | .050 | .061 | .022 | .64 | -.073 |
| Boundary Extension | .045 | -.034 | -.023 | -.035 | -.004 | .11 | .86 |
| **Variance explained (Total = 59.75%)** | **12.47** | **11.04** | **9.73** | **9.09** | **6.98** | **5.37** | **5.07** |

**Table S8.**
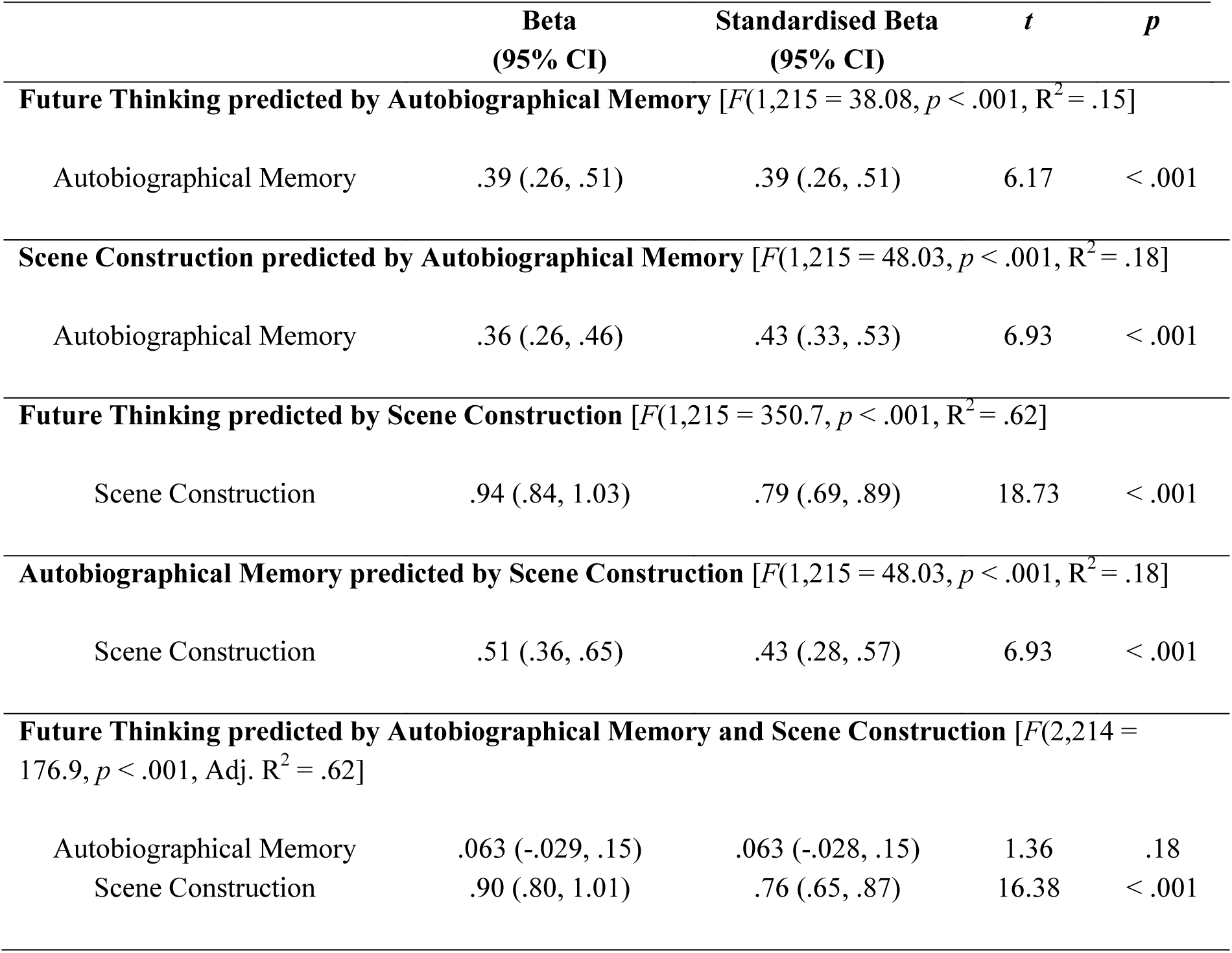
Full details of the regression analyses shown in Figure 1 examining the mediation analyses of the Scene component variables when future thinking is the dependent variable

**Table S9.** Full details of the regression analyses shown in Figure 3 examining the mediation analyses of the Scene component variables when future thinking is the independent variable

|  | **Beta**<br>**(95% CI)** | **Standardised Beta**<br>**(95% CI)** | ***t*** | ***p*** |
| --- | --- | --- | --- | --- |
| **Autobiographical Memory predicted by Future Thinking** [$F(1,215 = 38.08, p < .001, R^2 = .15]$] | | | | |
| Future Thinking | .39 (.26, .51) | .39 (.26,.51) | 6.17 | < .001 |
| **Scene Construction predicted by Future Thinking** [$F(1,215 = 350.7, p < .001, R^2 = .62]$] | | | | |
| Future Thinking | .66 (.59, .73) | .79 (.72, .86) | 18.73 | < .001 |
| **Autobiographical Memory predicted by Future Thinking and Scene Construction** [$F(2,214 = 25.03, p < .001, \text{Adj. } R^2 = .18]$] | | | | |
| Future thinking | .14 (-.062, .33) | .14 (-.062, .33) | 1.36 | .18 |
| Scene Construction | .38 (.15, .62) | .32 (.086, .56) | 3.21 | .0015 |
| **Scene Construction predicted by Future Thinking and Autobiographical Memory** [$F(2,214 = 188.1, p < .001, \text{Adj. } R^2 = .63]$] | | | | |
| Future thinking | .62 (.54, .69) | .73 (.66, .81) | 16.38 | < .001 |
| Autobiographical Memory | .12 (.047, .19) | .14 (.070, .22) | 3.21 | .0015 |

**Table S10.** Full details of the regression analyses shown in Figure 5 examining the mediation analyses of the scene construction, autobiographical memory and navigation relationships

|  | **Beta**<br>**(95% CI)** | **Standardised Beta**<br>**(95% CI)** | ***t*** | ***p*** |
| --- | --- | --- | --- | --- |
| **Navigation predicted by Autobiographical Memory** [$F(1,215) = 8.92, p = .0031, R^2 = .040$] | | | | |
| Autobiographical memory | .99 (.34, 1.64) | .20 (-.45, .85) | 2.99 | .0032 |
| **Scene Construction predicted by Autobiographical Memory** [$F(1,215) = 48.03, p < .001, R^2 = .18$] | | | | |
| Autobiographical Memory | .36 (.26, .46) | .43 (.33, .53) | 6.93 | <.001 |
| **Navigation predicted by Scene construction** [$F(1,215) = 19.83, p < .001, R^2 = .084$] | | | | |
| Scene Construction | 1.71 (.95, 2.47) | .29 (-.47, 1.05) | 4.45 | <.001 |
| **Autobiographical Memory predicted by Scene Construction** [$F(1,215) = 48.03, p < .001, R^2 = .18$] | | | | |
| Scene Construction | .51 (.36, .65) | .43 (.28, .57) | 6.93 | <.001 |
| **Navigation predicted by Autobiographical Memory and Scene Construction** [$F(2,214) = 10.76, p < .001, \text{Adj. } R^2 = .083$] | | | | |
| Autobiographical Memory | .46 (-.25, 1.16) | .092 (-.61, .79) | 1.28 | .20 |
| Scene Construction | 1.48 (.64, 2.32) | .25 (-.59, 1.09) | 3.49 | <.001 |

**Table S11.** Full details of the regression analyses shown in Figure 7 examining the mediation analyses of the future thinking to navigation relationship with scene construction or autobiographical memory as the mediating variable

|  | **Beta**<br>**(95% CI)** | **Standardised Beta**<br>**(95% CI)** | ***t*** | ***p*** |
| --- | --- | --- | --- | --- |
| **Navigation predicted by Future thinking** [$F(1,215 = 14.48, p < .001, R^2 = .063]$] | | | | |
| Future Thinking | 1.24 (.60, 1.89) | .25 (-.39, .90) | 3.81 | < .001 |
| **Scene Construction predicted by Future Thinking** [$F(1,215 = 350.7, p < .001, R^2 = .62]$] | | | | |
| Future Thinking | .66 (.59, .73) | .79 (.72, .86) | 18.73 | < .001 |
| **Autobiographical Memory predicted by Future Thinking** [$F(1,215 = 38.08, p < .001, R^2 = .15]$] | | | | |
| Future Thinking | .39 (.26, .51) | .39 (.26, .51) | 6.17 | < .001 |
| **Navigation predicted by Future Thinking and Scene Construction** [$F(2,214 = 10.04, p < .001, \text{Adj. } R^2 = .077]$] | | | | |
| Future thinking | .29 (-.74, 1.33) | .059 (-.98, 1.09) | .56 | .58 |
| Scene Construction | 1.44 (.21, 2.67) | .24 (-.99, 1.48) | 2.30 | .022 |
| **Navigation predicted by Future thinking and Autobiographical Memory** [$F(2,214 = 8.72, p < .001, \text{Adj. } R^2 = .067]$] | | | | |
| Future Thinking | 1.01 (.32, 1.71) | .20 (-.49, .90) | 2.87 | .0045 |
| Autobiographical Memory | .59 (-.10, 1.29) | .12 (-.57, .82) | 1.69 | .093 |

**Table S12.** Details of the pathways within the structural equation model of the Spatial component to navigation relationship, with the Scene component as the mediating variable

|  | **Beta**<br>**(95% CI)** | **Standardised Beta**<br>**(95% CI)** | **z** | **p** |
| --- | --- | --- | --- | --- |
| **Spatial Component latent variable predictors** |  |  |  |  |
| ROCF delayed recall | 3.87 (3.09, 4.66) | .68 (.58, .78) | 9.67 | < .001 |
| Paper Folding | 2.59 (2.06, 3.11) | .68 (.57, .78) | 9.61 | < .001 |
| Object-Place Association | 1.18 (.87, 1.48) | .55 (.44, .67) | 7.62 | < .001 |
| Brixton Spatial Anticipation Test | .66 (.38, .93) | .36 (.22, .49) | 4.71 | < .001 |
| **Scene Component latent variable predictors** |  |  |  |  |
| Scene Construction | 5.45 (4.67, 6.21) | .93 (.85, 1.01) | 14.05 | < .001 |
| Future Thinking | 5.87 (4.98, 6.76) | .84 (.77, .92) | 12.90 | < .001 |
| Autobiographical Memory | 3.21 (2.28, 4.14) | .46 (.35, .57) | 6.77 | < .001 |
| **Scene Component predicted by the Spatial Component** |  |  |  |  |
| Spatial Component | .28 (.1, .46) | .27 (.11, .43) | 3.06 | .002 |
| **Predictors of Navigation** |  |  |  |  |
| Spatial Component | 22.71 (17.64, 27.79) | .64 (.52, .75) | 8.77 | < .001 |
| Scene Component | 4.87 (.47, 9.27) | .14 (.016, .27) | 2.17 | .030 |

**Table S13.**
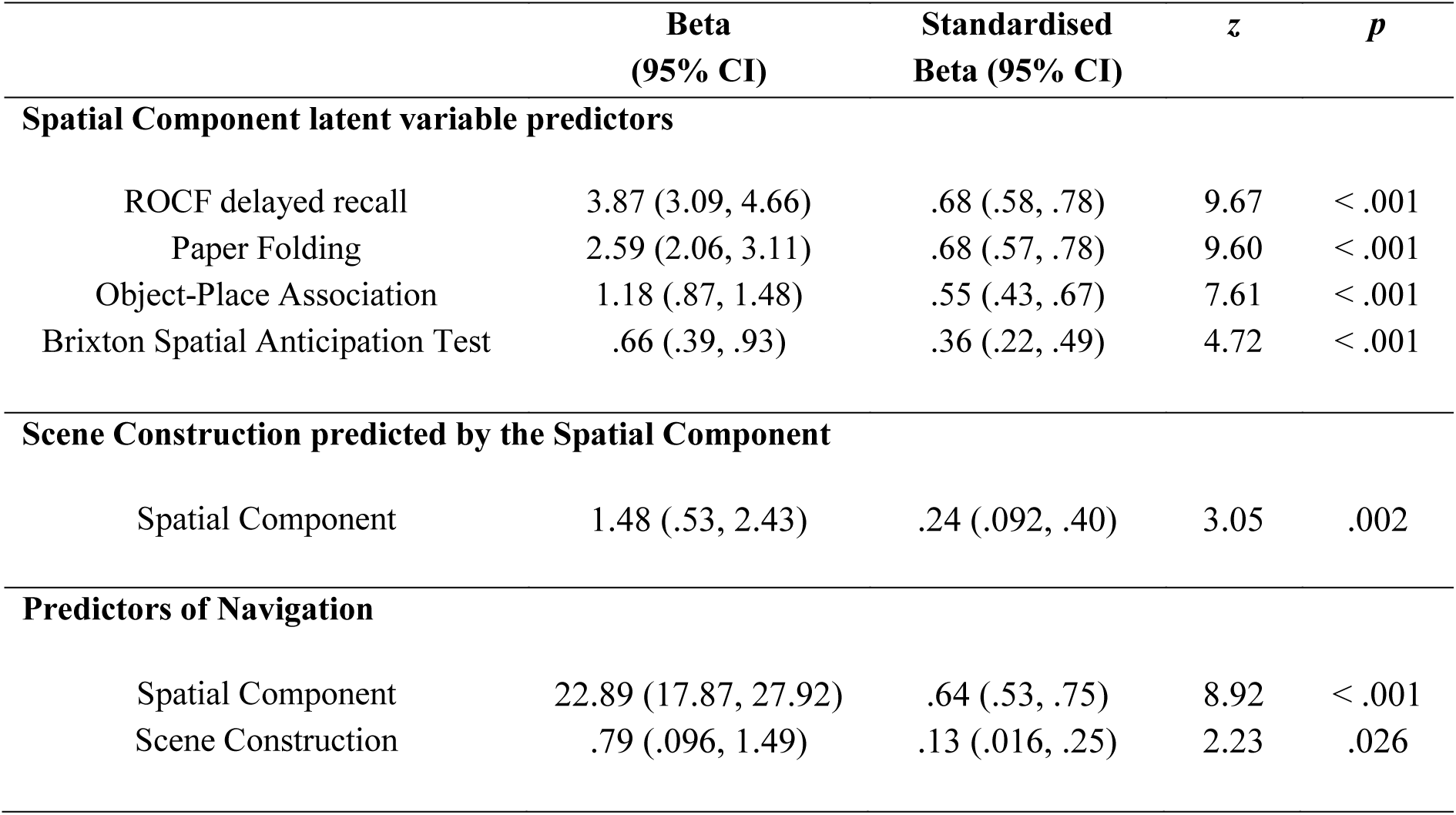
Details of the pathways within the structural equation model of the Spatial component to navigation relationship, with scene construction as the mediating variable

|  | **Beta**<br>**(95% CI)** | **Standardised**<br>**Beta (95% CI)** | **z** | **p** |
| --- | --- | --- | --- | --- |
| **Spatial Component latent variable predictors** |  |  |  |  |
| ROCF delayed recall | 3.87 (3.09, 4.66) | .68 (.58, .78) | 9.67 | < .001 |
| Paper Folding | 2.59 (2.06, 3.11) | .68 (.57, .78) | 9.60 | < .001 |
| Object-Place Association | 1.18 (.87, 1.48) | .55 (.43, .67) | 7.61 | < .001 |
| Brixton Spatial Anticipation Test | .66 (.39, .93) | .36 (.22, .49) | 4.72 | < .001 |
| **Scene Construction predicted by the Spatial Component** |  |  |  |  |
| Spatial Component | 1.48 (.53, 2.43) | .24 (.092, .40) | 3.05 | .002 |
| **Predictors of Navigation** |  |  |  |  |
| Spatial Component | 22.89 (17.87, 27.92) | .64 (.53, .75) | 8.92 | < .001 |
| Scene Construction | .79 (.096, 1.49) | .13 (.016, .25) | 2.23 | .026 |

**Table S14.** Details of the pathways within the structural equation model of the Spatial component to navigation relationship, with autobiographical memory as the mediating variable

|  | **Beta**<br>**(95% CI)** | **Standardised Beta**<br>**(95% CI)** | **z** | **p** |
| --- | --- | --- | --- | --- |
| **Spatial Component latent variable predictors** |  |  |  |  |
| ROCF delayed recall | 3.86 (3.08, 4.65) | .68 (.58, .78) | 9.63 | < .001 |
| Paper Folding | 2.59 (2.06, 3.12) | .68 (.57, .78) | 9.61 | < .001 |
| Object-Place Association | 1.17 (.87, 1.48) | .55 (.43, .67) | 7.60 | < .001 |
| Brixton Spatial Anticipation Test | .67 (.39, .94) | .36 (.22, .50) | 4.77 | < .001 |
| **Autobiographical Memory predicted by the Spatial Component** |  |  |  |  |
| Spatial Component | .81 (-.34, 1.96) | .11 (-.046, .27) | 1.38 | .17 |
| **Predictors of Navigation** |  |  |  |  |
| Spatial Component | 23.57 (18.67, 28.48) | .66 (.55, .76) | 9.42 | < .001 |
| Autobiographical Memory | .62 (.057, 1.19) | .13 (.012, .24) | 2.16 | .031 |

**Table S15.** Details of the pathways within the structural equation model of the Spatial component to navigation relationship, with future thinking as the mediating variable

|  | **Beta**<br>**(95% CI)** | **Standardised Beta**<br>**(95% CI)** | **z** | **p** |
| --- | --- | --- | --- | --- |
| **Spatial Component latent variable predictors** |  |  |  |  |
| ROCF delayed recall | 3.87 (3.08, 4.65) | .68 (.58, .78) | 9.65 | < .001 |
| Paper Folding | 2.59 (2.06, 3.11) | .68 (.57, .78) | 9.61 | < .001 |
| Object-Place Association | 1.18 (.88, 1.48) | .55 (.44, .67) | 7.65 | < .001 |
| Brixton Spatial Anticipation Test | .66 (.38, .93) | .36 (.22, .49) | 4.71 | < .001 |
| **Future Thinking predicted by the Spatial Component** |  |  |  |  |
| Spatial Component | 1.74 (.61, 2.87) | .24 (.089, .39) | 3.01 | .003 |
| **Predictors of Navigation** |  |  |  |  |
| Spatial Component | 23.26 (18.18, 28.33) | .65 (.54, .76) | 8.98 | < .001 |
| Future Thinking | .47 (-.12, 1.06) | .094 (-.025, .21) | 1.55 | .12 |

#### Control mediation analyses between the Verbal Memory component and the tasks of the Scene component

The Scene component of the PCA contained tasks that were scored from open ended verbal descriptions. As such, verbal task demands - be that narrative style, verbal ability and so forth - or similarities in scoring across the tasks could be candidate processes linking scene construction, autobiographical memory and future thinking. As we detail in the main text, we do not believe this to be the case due to the pattern of results that emerged. However, to further examine the potential involvement of verbal processing, we also ran a series of control mediation analyses looking at the effects of the Verbal Memory component (as a proxy for verbal ability) on the tasks of the Scene component.

We did this by employing the same methodology as when relating the Spatial and Scene components to navigation. In short, using SEM, a latent variable was used to represent the Verbal Memory component. The Verbal Memory latent variable was comprised of the tasks identified by the PCA, namely: Concrete Verbal Paired Associates, WMS Verbal Paired Associates, RAVLT, Abstract Verbal Paired Associates and the Logical Memory test.

Figure S2 shows the SEMs of the relationships between the Verbal Memory component and each of autobiographical memory, future thinking and scene construction when mediated by the other tasks of the Scene component (i.e. scene construction, autobiographical memory or future thinking). The latent variable (Verbal Memory) is shown in a circle, the observed variables (the cognitive tasks) in rectangles. The numerical values represent the standardised coefficients of the path in question. For all models, overall model fit was good, in line with published recommendations [a: *χ*^2^(13) = 18.52, *p* = .14; CFI = .98; TLI = .97; RMSEA = .044 (90% CI: 0, .086); SRMR = .037; b: *χ*^2^(13) = 14.23, *p* = .36; CFI = .997; TLI = .996; RMSEA = .021 (90% CI: 0, .072); SRMR = .029; c: *χ*^2^(13) = 18.52, *p* = .14; CFI = .98; TLI = .97; RMSEA = .044 (90% CI: 0, .086); SRMR = .037; d: *χ*^2^(13) = 18.14, *p* = .15; CFI = .98; TLI = .97; RMSEA = .043 (90% CI: 0, .085); SRMR = .036; e: *χ*^2^(13) = 14.23, *p* = .36; CFI = .997; TLI = .996; RMSEA = .021 (90% CI: 0, .072); SRMR = .029; f: *χ*^2^(13) = 18.14, *p* = .15; CFI = .98; TLI = .97; RMSEA = .043 (90% CI: 0, .085); SRMR = .036].

Of key relevance to our question, the influence of the Verbal Memory component was either fully or partially mediated in all the models. This suggests that the results reported in the main text showing the relationships between scene construction, autobiographical memory and future thinking cannot simply be explained by verbal ability.

**Figure S2.**
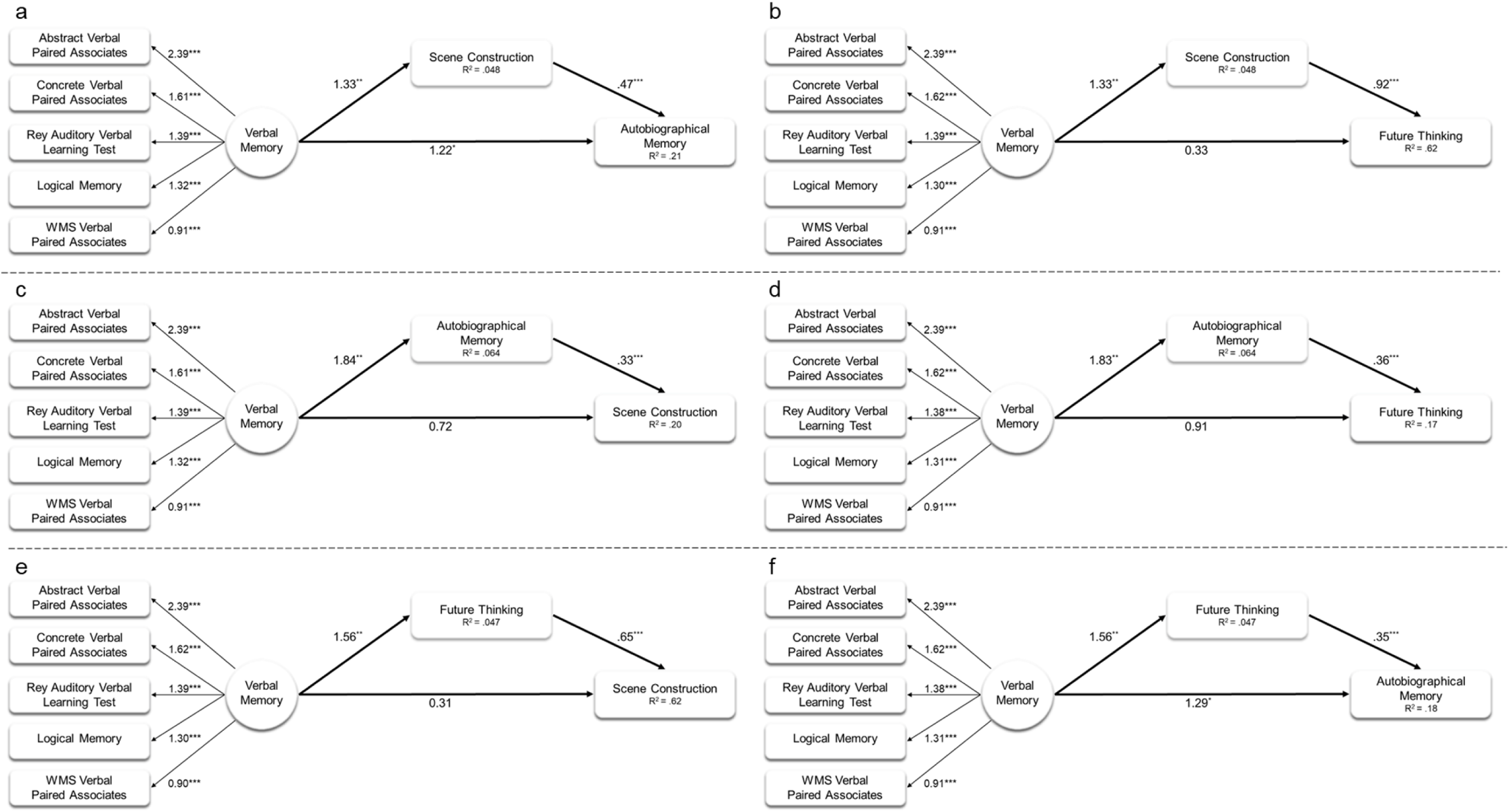
Structural equation models of the mediation effects of scene construction, autobiographical memory or future thinking on the Verbal Memory component to scene construction, autobiographical memory or future thinking relationship. The darker arrows show the main paths of interest, the lighter arrows show the links between the individual observed variables and the latent variable (Verbal Memory). The R2 values represent the proportion of variance explained by the main paths of interest (i.e. the dark arrows). Numerical values linked with a pathway represent standardized path coefficients. *p < .05, **p < .01, ***p < .001.

#### Control mediation analyses using the delayed recall of the ROCF as the mediator variable between the tasks of the Scene component

A potential criticism of our previous analyses is that we tested the influence of memory on the relationship between scene construction, autobiographical memory and future thinking using a memory task that relied on verbal output. To address this issue, we also examined the relationships between scene construction, autobiographical memory and future thinking with a nonverbal memory task (the delayed recall of the ROCF) as the mediator.

First, we ascertained whether mediation by the ROCF was possible. As detailed in the main text, according to the steps outlined by Baron & Kenny (1986), for mediation to occur, the independent variable must predict the mediator variable. However, while scene construction and future thinking predicted the ROCF, autobiographical memory did not (see Table S16). As such, while it has also been suggested that an initial direct relationship between the independent variable (autobiographical memory) and the dependent variable (ROCF) is not required in the presence of a strong a priori belief of a small effect size (e.g. Shrout & Bolger, 2002), given the substantial sample size of the current study and the observed effect size of 0.0047 we deemed that the ROCF could not, mediate the relationships between autobiographical memory and scene construction, and autobiographical memory and future thinking. Moreover, as no relationship was found between the ROCF and autobiographical memory, the ROCF could also not mediate between scene construction and autobiographical memory, or future thinking and autobiographical memory. As such, there were only two possible relationships that the ROCF could theoretically mediate – the scene construction to future thinking relationship, and the future thinking to scene construction relationship. However, mediation analyses found no effect of the ROCF on either (Tables S16–17). The ROCF did not, therefore, mediate between any of the tasks of the Scene component.

**Table S16.** Full details of the regression analyses examining the mediation analyses using the delayed recall of the ROCF as the mediator variable between the tasks of the Scene component

|  | **Beta**<br>**(95% CI)** | **Standardised Beta**<br>**(95% CI)** | ***t*** | ***p*** |
| --- | --- | --- | --- | --- |
| **ROCF delayed recall predicted by Scene Construction** [$F(1,215) = 8.67, p = .0036, R^2 = .039$] | | | | |
| Scene Construction | .18 (.061, .31) | .20 (.073, .32) | 2.95 | .0036 |
| **ROCF delayed recall predicted by Autobiographical Memory** [$F(1,215) = 1.02, p = .31, R^2 = .0047$] | | | | |
| Autobiographical Memory | .054 (-.05, .16) | .69 (-.38, .17) | 1.09 | .32 |
| **ROCF delayed recall predicted by Future Thinking** [$F(1,215) = 6.10, p = .014, R^2 = .023$] | | | | |
| Future Thinking | .13 (.026, .24) | .17 (.061, .27) | 2.47 | .014 |
| **Scene Construction predicted by Future Thinking and ROCF delayed recall** [$F(2,214) = 177.9, p < .001, \text{Adj. } R^2 = .62$] | | | | |
| Future thinking | .65 (-.74, 1.33) | .78 (.71, .85) | 18.27 | < .001 |
| Rey Figure | .072 (-.017, .16) | .067 (-.021, .16) | 1.60 | .11 |
| **Future Thinking predicted by Scene Construction and ROCF delayed recall** [$F(2,214) = 174.6, p < .001, \text{Adj. } R^2 = .62$] | | | | |
| Scene Construction | .93 (.83, 1.03) | .79 (.68, .89) | 18.27 | < .001 |
| Rey Figure | .015 (-.093, .12) | .011 (-.096, .12) | .27 | .79 |

**Table S17.**
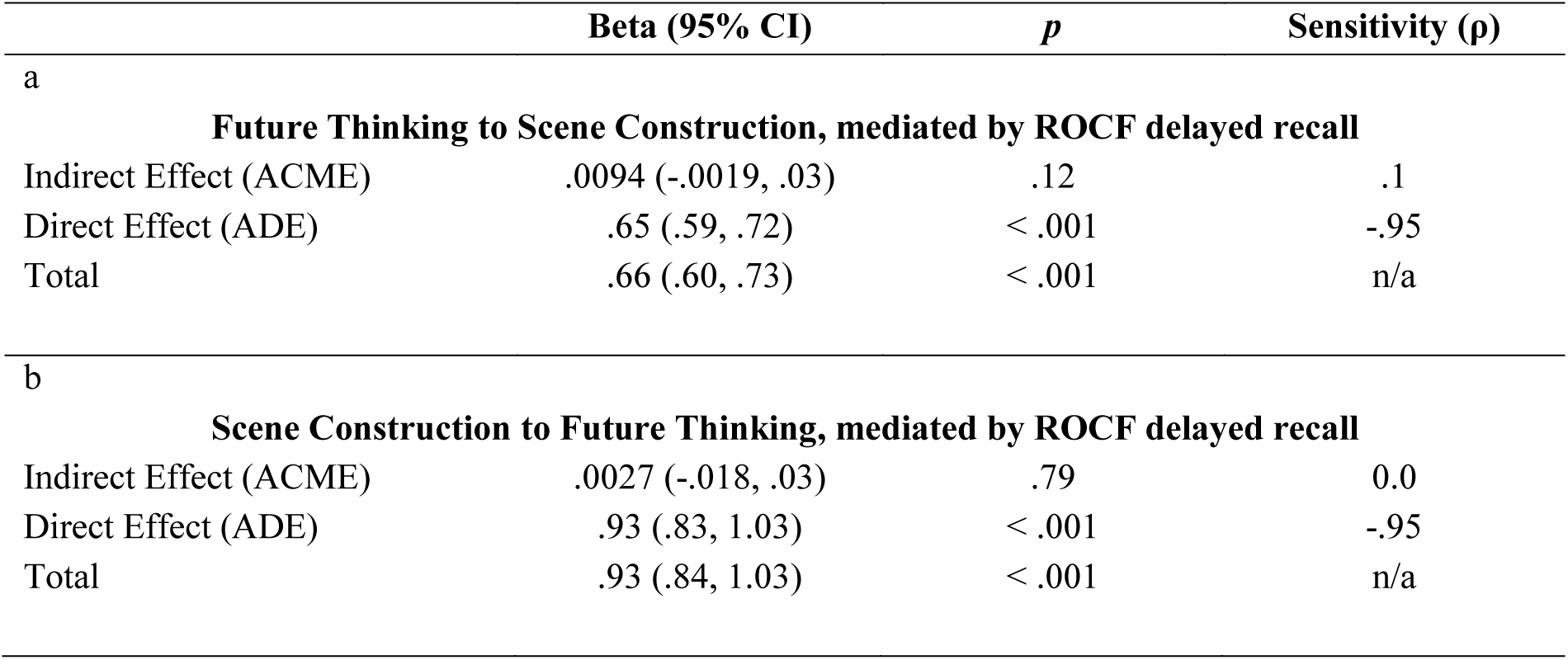
Mediation analyses of the Scene component variables with the delayed recall of the ROCF as the mediator variable

## References

Addis, D. R., Cheng, T., Roberts, R. P., & Schacter, D. L. (2011). Hippocampal contributions to the episodic simulation of specific and general future events. Hippocampus, 21(10), 1045–1052. doi: http://dx.doi.org/doi:10.1002/hipo.20870

Addis, D. R., Wong, A. T., & Schacter, D. L. (2007). Remembering the past and imagining the future: Common and distinct neural substrates during event construction and elaboration. Neuropsychologia, 45(7), 1363–1377. doi: http://dx.doi.org/10.1016/j.neuropsychologia.2006.10.016

Andelman, F., Hoofien, D., Goldberg, I., Aizenstein, O., & Neufeld, M. Y. (2010). Bilateral hippocampal lesion and a selective impairment of the ability for mental time travel. Neurocase, 16(5), 426–435. doi: http://dx.doi.org/10.1080/13554791003623318

Anderson, J. C., & Gerbing, D. W. (1988). Structural equation modeling in practice: A review and recommended two-step approach. Psychological Bulletin, 103(3), 411. doi: http://dx.doi.org/10.1037/0033-2909.103.3.411

Andrews-Hanna, J. R., Reidler, J. S., Sepulcre, J., Poulin, R., & Buckner, R. L. (2010). Functional-anatomic fractionation of the brain's default network. Neuron, 65(4), 550–562. doi: http://doi.org/10.1016/j.neuron.2010.02.005

Andrews-Hanna, J. R., Saxe, R., & Yarkoni, T. (2014). Contributions of episodic retrieval and mentalizing to autobiographical thought: Evidence from functional neuroimaging, resting-state connectivity, and fMRI meta-analyses. Neuroimage, 91, 324–335. doi: http://dx.doi.org/10.1016/j.neuroimage.2014.01.032

Arnold, K. M., McDermott, K. B., & Szpunar, K. K. (2011). Imagining the near and far future: The role of location familiarity. Memory and Cognition, 39(6), 954–967. doi: http://dx.doi.org/10.3758/s13421-011-0076-1

Baron, R. M., & Kenny, D. A. (1986). The moderator-mediator variable distinction in social psychological research: Conceptual, strategic, and statistical considerations. Journal of Personality and Social Psychology & Marketing, 51(6), 1173–1182. doi: http://dx.doi.org/10.1037//0022-3514.51.6.1173

Buckner, R. L., & Carroll, D. C. (2007). Self-projection and the brain. Trends in Cognitive Sciences, 11(2), 49–57. doi: http://dx.doi.org/10.1016/j.tics.2006.11.004

Burgess, P. W., & Shallice, T. (1997). The Hayling and Brixton Tests. Thurston, Suffolk: Thames Valley Test Company.

Cabeza, R., & St. Jacques, P. (2007). Functional neuroimaging of autobiographical memory. Trends in Cognitive Sciences, 11(5), 49–57. doi: http://dx.doi.org/10.1016/j.tics.2007.02.005

Cattell, R. B. (1978). The scientific use of factor analysis in behavioral and life sciences. New York: Plenum.

Ciaramelli, E., Faggi, G., Scarpazza, C., Mattioli, F., Spaniol, J., Ghetti, S., & Moscovitch, M. (2017). Subjective recollection independent from multifeatural context retrieval following damage to the posterior parietal cortex. Cortex, 91, 114–125. doi: http://dx.doi.org/10.1016/j.cortex.2017.03.015

Cipolotti, L., & Maguire, E. A. (2003). A combined neuropsychological and neuroimaging study of topographical and non-verbal memory in semantic dementia. Neuropsychologia, 41(9), 1148–1159. doi: http://dx.doi.org/10.1016/S0028-3932(03)00032-0

Clark, I. A., Kim, M., & Maguire, E. A. (2018). Verbal paired associates and the hippocampus: The role of scenes. Journal of Cogntive Neuroscience. doi: http://dx.doi.org/10.1101/206250

Clark, I. A., & Maguire, E. A. (2016). Remembering preservation in hippocampal amnesia. Annual Review of Psychology, 67(1), 51–82. doi: http://dx.doi.org/10.1146/annurev-psych-122414-033739

Cohen, J. (1992). A power primer. Psychological Bulletin, 112(1), 155–159. doi: http://dx.doi.org/10.1037/0033-2909.112.1.155

D’Argembeau, A., & Van der Linden, M. (2004). Phenomenal characteristics associated with projecting oneself back into the past and forward into the future: Influence of valence and temporal distance. Consciousness and Cognition, 13(4), 844–858. doi: http://dx.doi.org/10.1016/j.concog.2004.07.007

D’Argembeau, A., & Van der Linden, M. (2006). Individual differences in the phenomenology of mental time travel: The effect of vivid visual imagery and emotion regulation strategies. Consciousness and Cognition, 15(2), 342–350. doi: http://dx.doi.org/10.1016/j.concog.2005.09.001

Dalton, M. A., & Maguire, E. A. (2017). The pre/parasubiculum: a hippocampal hub for scene-based cognition? Current Opinion in Behavioral Sciences, 17, 34–40. doi: http://dx.doi.org/10.1016/j.cobeha.2017.06.001

Dalton, M. A., Zeidman, P., McCormick, C., & Maguire, E. A. (2018). Differentiable processing of objects, associations and scenes within the hippocampus. Journal of Neuroscience, 38(38), 8146–8159. doi: http://dx.doi.org/10.1523/JNEUROSCI.0263-18.2018

de Vito, S., Gamboz, N., & Brandimonte, M. A. (2012). What differentiates episodic future thinking from complex scene imagery? Consciousness and Cognition, 21(2), 813–823. doi: http://dx.doi.org/10.1016/j.concog.2012.01.013

Dimsdale-Zucker, H. R., Ritchey, M., Ekstrom, A. D., Yonelinas, A. P., & Ranganath, C. (2018). CA1 and CA3 differentially support spontaneous retrieval of episodic contexts within human hippocampal subfields. Nature Communications, 9(1), 294. doi: http://dx.doi.org/10.1038/s41467-017-02752-1

Eichenbaum, H., Yonelinas, A. R., & Ranganath, C. (2007). The medial temporal lobe and recognition memory. Annual Review of Neuroscience, 30, 123–152. doi: http://dx.doi.org/10.1146/annurev.neuro.30.051606.094328

Ekstrom, A. D., Kahana, M. J., Caplan, J. B., Fields, T. A., Isham, E. A., Newman, E. L., & Fried, I. (2003). Cellular networks underlying human spatial navigation. Nature, 425(6954), 184–188. doi: http://www.nature.com/nature/journal/v425/n6954/suppinfo/nature01964_S1.html

Ekstrom, R. B., French, J. W., Harman, H. H., & Dermen, D. (1976). Manual for kit of factor referenced cognitive tests: Educational Testing Service.

Epstein, R. A., Patai, E. Z., Julian, J. B., & Spiers, H. J. (2017). The cognitive map in humans: spatial navigation and beyond. Nature Neuroscience, 20, 1504. doi: http://dx.doi.org/10.1038/nn.4656

Fabrigar, L. R., Wegener, D. T., MacCallum, R. C., & Strahan, E. J. (1999). Evaluating the use of exploratory factor analysis in psychological research. Psychological Methods, 4(3), 272. doi: http://dx.doi.org/10.1037//1082-989x.4.3.272

Fava, J. L., & Velicer, W. F. (1992). The Effects of Overextraction on Factor and Component Analysis. Multivariate behavioral research, 27(3), 387–415. doi: http://dx.xoi.org/10.1207/s15327906mbr2703_5

Graham, K. S., Barense, M. D., & Lee, A. C. H. (2010). Going beyond LTM in the MTL: A synthesis of neuropsychological and neuroimaging findings on the role of the medial temporal lobe in memory and perception. Neuropsychologia, 48(4), 831–853. doi: http://dx.doi.org/10.1016/j.neuropsychologia.2010.01.001

Greenberg, D. L., & Knowlton, B. J. (2014). The role of visual imagery in autobiographical memory. Memory and Cognition, 42(6), 922–934. doi: http://dx.doi.org/10.3758/s13421-014-0402-5

Hassabis, D., Kumaran, D., & Maguire, E. A. (2007). Using imagination to understand the neural basis of episodic memory. Journal of Neuroscience, 27(52), 14365–14374. doi: http://dx.doi.org/10.1523/jneurosci.4549-07.2007

Hassabis, D., Kumaran, D., Vann, S. D., & Maguire, E. A. (2007). Patients with hippocampal amnesia cannot imagine new experiences. Proceedings of the National Academy of Sciences, 104(5), 1726–1731. doi: http://dx.doi.org/10.1073/pnas.0610561104

Hassabis, D., & Maguire, E. A. (2007). Deconstructing episodic memory with construction. Trends in Cognitive Sciences, 11(7), 299–306. doi: http://dx.doi.org/10.1016/j.tics.2007.05.001

Hebscher, M., Levine, B., & Gilboa, A. (2017). The precuneus and hippocampus contribute to individual differences in the unfolding of spatial representations during episodic autobiographical memory. Neuropsychologia. doi: http://dx.doi.org/10.1016/j.neuropsychologia.2017.03.029

Hodgetts, C. J., Voets, N. L., Thomas, A. G., Clare, S., Lawrence, A. D., & Graham, K. S. (2017). Ultra-high-field fMRI reveals a role for the subiculum in scene perceptual discrimination. Journal of Neuroscience, 37(12), 3150–3159. doi: http://dx.doi.org/10.1523/JNEUROSCI.3225-16.2017

Hu, L., & Bentler, P. M. (1999). Cutoff criteria for fit indexes in covariance structure analysis: Conventional criteria versus new alternatives. Structural Equation Modeling: A Multidisciplinary Journal, 6(1), 1–55. doi: http://dx.doi.org/10.1080/10705519909540118

Imai, K., Keele, L., & Tingley, D. (2010). A general approach to causal mediation analysis. Psychological Methods, 15(4), 309–334. doi: http://dx.doi.org/10.1037/a0020761

Intraub, H., & Richardson, M. (1989). Wide-angle memories of close-up scenes. Journal of Experimental Psychology: Learning, Memory, and Cognition, 15(2), 179–187. doi: http://dx.doi.org/10.1037/0278-7393.15.2.179

Kapur, N., Young, A., Bateman, D., & Kennedy, P. (1989). Focal retrograde amnesia: A long term clinical and neuropsychological follow-up. Cortex, 25(3), 387–402. doi: http://dx.doi.org/10.1016/S0010-9452(89)80053-X

Klein, S. B., Loftus, J., & Kihlstrom, J. F. (2002). Memory and temporal experience: The effects of episodic memory loss on an amnesic patient's ability to remember the past and imagine the future. Social Cognition, 20(5), 353–379. doi: http://dx.doi.org/10.1521/soco.20.5.353.21125

Konkle, T., Brady, T. F., Alvarez, G. A., & Oliva, A. (2010). Scene memory is more detailed than you think:The role of categories in visual long-term memory. Psychological Science, 21(11), 1551–1556. doi: http://dx.doi.org/10.1177/0956797610385359

Kozhevnikov, M., Kosslyn, S., & Shephard, J. (2005). Spatial versus object visualizers: A new characterization of visual cognitive style. Memory and Cognition, 33(4), 710- 726. doi: http://dx.doi.org/10.3758/BF03195337

Kraemer, D. J., Schinazi, V. R., Cawkwell, P. B., Tekriwal, A., Epstein, R. A., & Thompson-Schill, S. L. (2017). Verbalizing, visualizing, and navigating: The effect of strategies on encoding a large-scale virtual environment. Journal of Experimental Psychology: Learning, Memory, and Cognition, 43(4), 611. doi: http://dx.doi.org/10.1037/xlm0000314

Lee, A. C. H., Bussey, T. J., Murray, E. A., Saksida, L. M., Epstein, R. A., Kapur, N., . . . Graham, K. S. (2005). Perceptual deficits in amnesia: challenging the medial temporal lobe ‘mnemonic’ view. Neuropsychologia, 43(1), 1–11. doi: http://dx.doi.org/10.1016/j.neuropsychologia.2004.07.017

Levine, B., Svoboda, E., Hay, J. F., Winocur, G., & Moscovitch, M. (2002). Aging and autobiographical memory: Dissociating episodic from semantic retrieval. Psychology and Aging, 17(4), 677–689. doi: http://dx.doi.org/10.1037/0882-7974.17.4.677

Maguire, E. A., Gadian, D. G., Johnsrude, I. S., Good, C. D., Ashburner, J., Frackowiak, R. S. J., & Frith, C. D. (2000). Navigation-related structural change in the hippocampi of taxi drivers. Proceedings of the National Academy of Sciences, 97(8), 4398–4403. doi: http://dx.doi.org/10.1073/pnas.070039597

Maguire, E. A., & Mullally, S. L. (2013). The hippocampus: A manifesto for change. Journal of Experimental Psychology: General, 142(4), 1180–1189. doi: http://dx.doi.org/10.1037/a0033650

McCormick, C., Rosenthal, C. R., Miller, T. D., & Maguire, E. A. (2017). Deciding what is possible and impossible following hippocampal damage in humans. Hippocampus, 27(3), 303–314. doi: http://dx.doi.org/10.1002/hipo.22694

Moffat, S. D., Elkins, W., & Resnick, S. M. (2006). Age differences in the neural systems supporting human allocentric spatial navigation. Neurobiology of Aging, 27(7), 965–972. doi: http://dx.doi.org/10.1016/j.neurobiolaging.2005.05.011

Moscovitch, M., Cabeza, R., Winocur, G., & Nadel, L. (2016). Episodic memory and beyond: The hippocampus and neocortex in transformation. Annual Review of Psychology, 67(1), 105–134. doi: http://dx.doi.org/doi:10.1146/annurev-psych-113011-143733

Moser, E. I., Kropff, E., & Moser, M.-B. (2008). Place cells, grid cells, and the brain's spatial representation system. Annual Review of Neuroscience, 31(1), 69–89. doi: http://dx.doi.org/10.1146/annurev.neuro.31.061307.090723

Mullally, S. L., Intraub, H., & Maguire, E. A. (2012). Attenuated boundary extension produces a paradoxical memory advantage in amnesic patients. Current Biology, 22(4), 261–268. doi: http://dx.doi.org/10.1016/j.cub.2012.01.001

O’Keefe, J., & Dostrovsky, J. (1971). The hippocampus as a spatial map. Preliminary evidence from unit activity in the freely-moving rat. Brain Research, 34(1), 171–175. doi: http://dx.doi.org/10.1016/0006-8993(71)90358-1

O’Keefe, J., & Nadel, L. (1978). The hippocampus as a cognitive map. Oxford: Clarendon Press.

Palombo, D. J., Hayes, S. M., Peterson, K. M., Keane, M. M., & Verfaellie, M. (2018). Medial temporal lobe contributions to episodic future thinking: Scene construction or future projection? Cerebral Cortex, 28(2), 447–458. doi: http://dx.doi.org/10.1093/cercor/bhw381

Race, E., Keane, M. M., & Verfaellie, M. (2011). Medial temporal lobe damage causes deficits in episodic memory and episodic future thinking not attributable to deficits in narrative construction. Journal of Neuroscience, 31(28), 10262–10269. doi: http://dx.doi.org/10.1523/jneurosci.1145-11.2011

Ramanan, S., Alaeddin, S., Goldberg, Z.-l., Strikwerda-Brown, C., Hodges, J. R., & Irish, M. (2018). Exploring the contribution of visual imagery to scene construction – Evidence from Posterior Cortical Atrophy. Cortex, 106, 261–274. doi: http://dx.doi.org/10.1016/j.cortex.2018.06.016

Rey, A. (1941). L'examen psychologique dans les cas d'encéphalopathie traumatique. (Les problems.). [The psychological examination in cases of traumatic encepholopathy. Problems.]. Archives de Psychologie, 28, 215–285.

Roberts, R. P., Schacter, D. L., & Addis, D. R. (2017). Scene construction and relational processing: Separable constructs? Cerebral Cortex, 1–4. doi: http://dx/.doi.org/10.1093/cercor/bhx081

Robin, J. (2018). Spatial scaffold effects in event memory and imagination. Wiley Interdisciplinary Reviews: Cognitive Science, e1462. doi: http://dx.doi.org/10.1002/wcs.1462

Robin, J., & Moscovitch, M. (2014). The effects of spatial contextual familiarity on remembered scenes, episodic memories, and imagined future events. Journal of Experimental Psychology: Learning, Memory, and Cognition, 40(2), 459–475. doi: http://dx.doi.org/10.1037/a0034886

Robin, J., Wynn, J., & Moscovitch, M. (2016). The spatial scaffold: The effects of spatial context on memory for events. Journal of Experimental Psychology: Learning, Memory, and Cognition, 42(2), 308–315. doi: http://dx.doi.org/10.1037/xlm0000167

Rosenbaum, R. S., Gilboa, A., Levine, B., Winocur, G., & Moscovitch, M. (2009). Amnesia as an impairment of detail generation and binding: Evidence from personal, fictional, and semantic narratives in K.C. Neuropsychologia, 47(11), 2181–2187. doi: http://dx.doi.org/10.1016/j.neuropsychologia.2008.11.028

Rosenbaum, R. S., Köhler, S., Schacter, D. L., Moscovitch, M., Westmacott, R., Black, S. E., . . . Tulving, E. (2005). The case of K.C.: Contributions of a memory-impaired person to memory theory. Neuropsychologia, 43(7), 989–1021. doi: http://dx.doi.org/10.1016/j.neuropsychologia.2004.10.007

Rosseel, Y. (2012). Lavaan: An R package for structural equation modeling. Journal of Statistical Software, 48(2), 1–36. doi: http://dx.doi.org/10.18637/jss.v048.i02

Rubin, D. C., & Umanath, S. (2015). Event memory: A theory of memory for laboratory, autobiographical, and fictional events. Psychological Review, 122(1), 1. doi: http://dx.doi.org/10.1037/a0037907

Rummel, R. J. (1970). Applied factor analysis. Evanston, IL: Northwestern University Press.

Schacter, D. L., Addis, D. R., & Buckner, R. L. (2007). Remembering the past to imagine the future: The prospective brain. Nature Reviews: Neuroscience, 8, 657–661. doi: http://dx.doi.org/10.1038/nrn2213

Schacter, D. L., Addis, D. R., Hassabis, D., Martin, V. C., Spreng, R. N., & Szpunar, K. K. (2012). The future of memory: Remembering, imagining, and the brain. Neuron, 76(4), 677–694. doi: http://dx.doi.org/10.1016/j.neuron.2012.11.001

Scoville, W. B., & Milner, B. (1957). Loss of recent memory after bilateral hippocampal lesions. Journal of Neurology, Neurosurgery and Psychiatry, 20, 11–21. doi: http://dx.doi.org/10.1136/jnnp.20.1.11

Sheldon, S., & Chu, S. (2017). What versus where: Investigating how autobiographical memory retrieval differs when accessed with thematic versus spatial information. The Quarterly Journal of Experimental Psychology, 70(9), 1909–1921. doi: http://dx.doi.org/10.1080/17470218.2016.1215478

Sheldon, S., & Levine, B. (2016). The role of the hippocampus in memory and mental construction. Annals of the New York Academy of Sciences, 1369(1), 76–92. doi: http://dx.doi.org/10.1111/nyas.13006

Shohamy, D., & Turk-Browne, N. B. (2013). Mechanisms for widespread hippocampal involvement in cognition. Journal of Experimental Psychology: General, 142(4), 1159–1170. doi: http://dx.doi.org/10.1037/a0034461

Shrout, P. E., & Bolger, N. (2002). Mediation in experimental and nonexperimental studies: new procedures and recommendations. Psychological Methods, 7(4), 422–445. doi: http://dx.doi.org/10.1037//1082-989X.7.4.422

Simons, J. S., Peers, P. V., Mazuz, Y. S., Berryhill, M. E., & Olson, I. R. (2010). Dissociation between memory accuracy and memory confidence following bilateral parietal lesions. Cerebral Cortex, 20(2), 479–485. doi: http://dx.doi.org/10.1093/cercor/bhp116

Smith, C. N., Jeneson, A., Frascino, J. C., Kirwan, C. B., Hopkins, R. O., & Squire, L. R. (2014). When recognition memory is independent of hippocampal function. Proceedings of the National Academy of Sciences, 111(27), 9935–9940. doi: http://dx.doi.org/10.1073/pnas.1409878111

Squire, L. R. (1992). Memory and the hippocampus: A synthesis from findings with rats, monkeys, and humans. Psychological Review, 99(2), 195–231. doi: http://dx.doi.org/10.1037/0033-295X.99.2.195

St-Laurent, M., Moscovitch, M., & McAndrews, M. P. (2016). The retrieval of perceptual memory details depends on right hippocampal integrity and activation. Cortex, 84, 15–33. doi: http://dx.doi.org/10.1016/j.cortex.2016.08.010

St. Jacques, P. L., Carpenter, A. C., Szpunar, K. K., & Schacter, D. L. (2018). Remembering and imagining alternative versions of the personal past. Neuropsychologia, 110, 170–179. doi: http://dx.doi.org/10.1016/j.neuropsychologia.2017.06.015

St. Jacques, P. L., Conway, M. A., Lowder, M. W., & Cabeza, R. (2010). Watching my mind unfold versus yours: An fMRI study using a novel camera technology to examine neural differences in self-projection of self versus other perspectives. Journal of Cognitive Neuroscience, 23(6), 1275–1284. doi: http://dx.doi.org/10.1162/jocn.2010.21518

Stawarczyk, D., & D'Argembeau, A. (2015). Neural correlates of personal goal processing during episodic future thinking and mind-wandering: An ALE meta-analysis. Human Brain Mapping, 36(8), 2928–2947. doi: http://dx.doi.org/10.1002/hbm.22818

Strauss, E., Sherman, E. M., & Spreen, O. (2006). A compendium of neuropsychological tests: administration, commentary and norms (3rd ed.). New York: Oxford University Press.

Svoboda, E., McKinnon, M. C., & Levine, B. (2006). The functional neuroanatomy of autobiographical memory: A meta-analysis. Neuropsychologia, 44(12), 2189–2208. doi: http://doi.org/10.1016/j.neuropsychologia.2006.05.023

Szpunar, K. K., & McDermott, K. B. (2008). Episodic future thought and its relation to remembering: Evidence from ratings of subjective experience. Consciousness and Cognition, 17(1), 330–334. doi: http://dx.doi.org/10.1016/j.concog.2007.04.006

Thakral, P. P., Benoit, R. G., & Schacter, D. L. (2017). Characterizing the role of the hippocampus during episodic simulation and encoding. Hippocampus, 27(12), 1275–1284. doi: http://dx.doi.org/doi:10.1002/hipo.22796

Tulving, E. (1985). Memory and consciousness. Canadian Psychology. Psychologie Canadienne, 26, 1–12. doi: http://dx.doi.org/10.1037/h0080017

Tulving, E. (2002). Episodic memory: From mind to brain. Annual Review of Psychology, 53(1), 1–25. doi: http://dx.doi.org/10.1146/annurev.psych.53.100901.135114

Verfaellie, M., & Keane, M. M. (2017). Neuropsychological investigations of human amnesia: Insights into the role of the medial temporal lobes in cognition. Journal of the International Neuropsychological Society, 23(9-10), 732–740. doi: http://dx.doi.org/10.1017/S1355617717000649

Warrington, E. K. (1984). Recognition Memory Test: Manual. Berkshire, UK: NFER-Nelson.

Wechsler, D. (2008). Wechsler adult intelligence scale–Fourth edition (WAIS–IV). San Antonio, TX: NCS Pearson.

Wechsler, D. (2009). WMS-IV.: Wechsler memory scale-Administration and scoring manual: Psychological Corporation.

Wechsler, D. (2011). Test of premorbid functioning. UK version (TOPF UK). London: Pearson Assessment.

Winocur, G., & Moscovitch, M. (2011). Memory transformation and systems consolidation. Journal of the International Neuropsychological Society, 17(5), 766–780. doi: http://dx.doi.org/10.1017/s1355617711000683

Wood, J. M., Tataryn, D. J., & Gorsuch, R. L. (1996). Effects of under-and overextraction on principal axis factor analysis with varimax rotation. Psychological Methods, 1(4), 354. doi: http://dx.doi.org/10.1037//1082-989x.1.4.354

Woollett, K., & Maguire, E. A. (2009). Navigational expertise may compromise anterograde associative memory. Neuropsychologia, 47(4), 1088–1095. doi: http://dx.doi.org/10.1016/j.neuropsychologia.2008.12.036

Woollett, K., & Maguire, E. A. (2010). The effect of navigational expertise on wayfinding in new environments. Journal of Environmental Psychology, 30(4), 565–573. doi: http://dx.doi.org/10.1016/j.jenvp.2010.03.003.

Zeidman, P., & Maguire, E. A. (2016). Anterior hippocampus: the anatomy of perception, imagination and episodic memory. Nature Reviews Neuroscience, 17(3), 173–182. doi: http://dx.doi.org/10.1038/nrn.2015.24

Zeidman, P., Lutti, A., & Maguire, E. A. (2015). Investigating the functions of subregions within anterior hippocampus. Cortex, 73, 240–256. doi: http://dx.doi.org/10.1016/j.cortex.2015.09.002

Zhong, J. Y., & Moffat, S. D. (2018). Extrahippocampal contributions to age-related changes in spatial navigation ability. Frontiers in Human Neuroscience, 12(272). doi: http://dx.doi.org/10.3389/fnhum.2018.00272

